# How Subjective Idea Valuation Energizes and Guides Creative Idea Generation

**DOI:** 10.1101/2022.08.02.502491

**Authors:** Alizée Lopez-Persem, Sarah Moreno Rodriguez, Marcela Ovando-Tellez, Théophile Bieth, Stella Guiet, Jules Brochard, Emmanuelle Volle

**Affiliations:** FrontLab, Sorbonne University, Institut du Cerveau - Paris Brain Institute - ICM, Inserm, CNRS, AP-HP, Hôpital de la Pitié Salpêtrière, Paris, France; Neurology department, Hôpital de la Pitié Salpêtrière, AP-HP, F-75013, Paris, France; TRA, “Life and Health”, University of Bonn, Bonn, Germany

**Author notes:** Corresponding authors: Alizée Lopez-Persem & Emmanuelle Volle. **Author Contributions:** EV, ALP designed the study. SMR, SG collected the data and performed model-free behavioral analyses. TB, MOT, SMR analysed the creativity battery data. ALP, EV, JB conceptualized the computational model. ALP performed the model-free and computational modelling analyses. ALP, EV wrote the article. All authors reviewed and edited the article. **Competing Interest Statement:** Authors declare no competing interests. **Data and Code Availability:** Data and scripts are currently published on gitlab. Access will be made open upon publication, or on request by editors or reviewers. **Author Node**.

**Keywords:** Creativity, preferences, computational modeling, subjective value, idea generation, evaluation

## Abstract

What drives us to search for creative ideas, and why does it feel good to find one? While previous studies demonstrated the positive influence of motivation on creative abilities, how reward and subjective values play a role in creativity remains unknown. This study proposes to characterize the role of individual preferences (how people value ideas) in creative ideation via behavioral experiments and computational modeling. Using the Free Generation of Associates Task coupled with rating tasks, we demonstrate the involvement of valuation processes during idea generation: preferred ideas are provided faster. We found that valuation depends on the adequacy and originality of ideas and guides response selection and creativity. Finally, our computational model correctly predicts the speed and quality of human creative responses, as well as interindividual differences in creative abilities. Altogether, this model introduces the mechanistic role of valuation in creativity. It paves the way for a neurocomputational account of creativity mechanisms.

**Public Significance Statement:** This study addresses the role of individual preferences in creativity. It demonstrates that preferences for ideas energize creative idea production: the more participants like their ideas, the faster they provide them. Moreover, preferences rely on an equilibrium between the adequacy and originality of ideas and vary across individuals. This study introduces a computational model which incorporates individual preferences and that correctly predicts the speed and quality of responses in a creative idea generation task, as well as inter-individual differences in creative abilities. Comparison of several versions of this model demonstrated that preferences guide the selection of creative responses.

---

Creativity is a core component of our ability to promote and cope with change. Creativity is defined as the ability to produce an object (or an idea) that is both original and adequate to the context (Dietrich, 2004; Runco & Jaeger, 2012; Jung & Vartanian, 2018). The cognitive mechanisms underlying the production of an original and adequate idea are yet to be elucidated.

It is largely admitted that creativity involves two interacting phases: generation and evaluation (Dietrich, 2004; Ellamil et al., 2012; Sowden et al., 2015; Beaty et al., 2016; Benedek & Jauk, 2018; Lin & Vartanian, 2018; Mekern et al., 2019; Kleinmintz et al., 2019; Guo et al., 2022). Theoretical models including these two processes have been proposed, such as the “two-fold model of creativity” (Kleinmintz et al., 2019), or the “blind-variation and selective retention model” (Campbell, 1960; Simonton, 1998; Sowden et al., 2015), a Darwinian-inspired theory stating that ideas are generated and evaluated on a trial and error basis, similarly to a variation-selection process. However, what kind of processes underlies evaluation in the context of creativity (in other words, what evaluative processes drive selection) remains overlooked.

Previous frameworks assumed that the originality and adequacy of ideas are evaluated to drive the selection of an idea during idea production (Donzallaz et al., 2021; Khalil & Moustafa, 2022; Lin & Vartanian, 2018). Existing theories also usually align evaluation with controlled or metacognitive processes (i.e., detecting relevant ideas, monitoring and applying some control to select or inhibit early thoughts and adapt to the context) and align them to the salience and executive control networks (ECN) (Beaty et al., 2014; Ellamil et al., 2012; Huang et al., 2015, 2018; Kleinmintz et al., 2019; Lin & Vartanian, 2018; Mayseless et al., 2014; Rataj et al., 2018; Ren et al., 2020; Rominger et al., 2020; Sowden et al., 2015). However, how these processes work and result in idea selection remains unknown. Because evaluative processes in other domains involve that subjective values are assigned to options to guide selection (Rushworth & Behrens, 2008), we hypothesize that evaluation in the context of creativity also requires building a subjective value. As previous work highlighted the importance of adequacy and originality in idea evaluation, we propose that this value is based on a combination of originality and adequacy of candidate ideas. Hence, we introduce valuation in the ideation process and dissociate them from other evaluation and generation processes. Valuation can be defined as a quantification of the subjective desire or preference for an entity (Redish et al., 2016) and consists in assigning a subjective value to an option, i.e., to define how much it is “likeable”, or “desirable”.

Previous studies assessing the role of evaluation in creativity (Ellamil et al., 2012; Huang et al., 2015, 2018; Mayseless et al., 2014; Rataj et al., 2018; Ren et al., 2020; Rominger et al., 2020) did not dissociate the valuation processes per se from the ones associated with controlled or metacognitive processes (i.e., evaluation, monitoring and applying some control to select or inhibit early thoughts and adapt to the context). However, the neuroscience of value-based decision-making demonstrated that they are distinct, experimentally dissociable, and have separate brain substrates (Shenhav & Karmarkar, 2019). Indeed, valuation processes have been investigated for centuries by philosophers, economists, psychologists, and more recently by neuroscientists (Levy & Glimcher, 2012), outside of the creativity field. Advances in the neuroscience of decision-making have allowed the identification of a neural network, the Brain Valuation System (BVS), representing the subjective value of options an agent considers (Levy & Glimcher, 2012). The BVS activity reflects values in a generic (independent of the kind of items) and automatic (even when we are engaged in another task) manner (Lopez-Persem et al., 2016). Interestingly, the BVS is often coupled with the ECN when a choice has to be made, in two different ways. First, in a top-down manner: the ECN modulates values according to the context (Hare et al., 2009); and second, in a bottom-up way: it drives the choice selection by integrating decision-values (Domenech et al., 2018). The new framework that we propose through the present study is that evaluation processes in creativity involve valuation, implemented by the BVS, in interaction with exploration and selection processes, supported by other networks.

Existing studies provide indirect arguments for the involvement of the BVS in creativity by showing a role of dopamine (Ang et al., 2018; Boot et al., 2017; Chermahini & Hommel, 2010; Manzano et al., 2010) and of the ventral striatum in creativity (Aberg et al., 2017; Huang et al., 2015; Takeuchi et al., 2010; Tik et al., 2018). Nevertheless, very little is known about the role of the BVS in creativity, and its interaction with the commonly reported brain networks for creativity (Default Mode Network (DMN) and ECN) has, to our knowledge, not been explored. In fact, the place for valuation processes in creativity still needs to be conceptualized and empirically investigated.

Here, we formulate the hypothesis that originality and adequacy are combined into a “subjective value” according to individual preferences, and that this subjective value drives the creative degree of the output. This value can impact the selection of an idea and possibly have a motivational role (Pessiglione et al., 2018) in exploring candidate ideas. Taking into account previous research from both creativity and decision-making fields, we hypothesize that creativity involves i) an *explorer* module that works on individual knowledge representations and provides a set of options/ideas varying in originality and adequacy; ii) a *valuator* module that computes the likeability of candidate ideas (their subjective value) based on a combination of their originality and adequacy with the goal an agent tries to reach; iii) a *selector* module that applies contextual constraints and integrates the subjective value of candidate ideas to guide the selection. To test these hypotheses, we combined several methods of cognitive and computational neuroscience. We built a computational model composed of the *explorer*, *valuator,* and *selector* modules, which we modeled separately (Figure 1) as detailed below.

**Figure 1.**
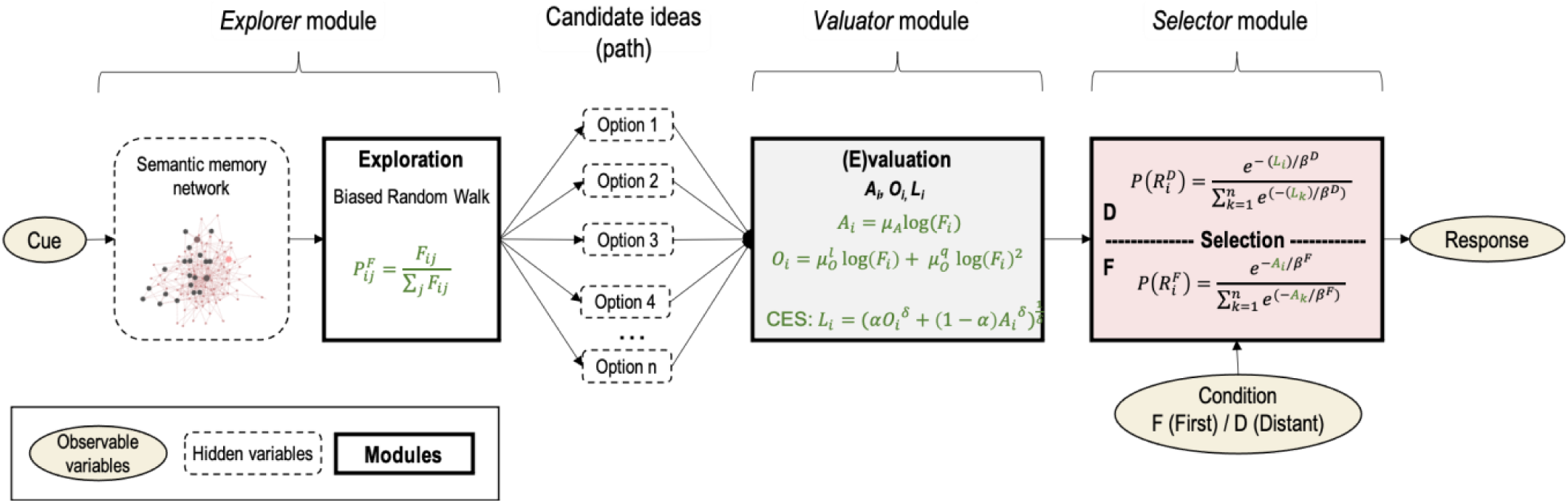
Schematic representation of the computational model. The model takes as input a cue, that “activates” a semantic memory network. Semantic search (exploration) is implemented as a biased random walk, in which node transition probability P is determined by the frequency of association F between the node i and its connected nodes j. The visited nodes (option 1 to n) are evaluated in terms of adequacy (A), originality (O) and the valuator assigns a likeability (L) to each of them, CES stands for Constant Elasticity of Substitution, see Results. A response is selected in function of the FGAT condition: in the *First* condition (F), the selection is based on adequacy and in the *Distant* condition (D), the selection is based on likeability. Equations results from the different model comparisons conducted in the study and are detailed in the manuscript. Text in black corresponds to our framework and hypotheses while text green corresponds to the results obtained in our study.

First, producing something new and appropriate (i.e., creative) relies in part on the ability to retrieve, manipulate or combine elements of knowledge stored in our memory (Benedek et al., 2012; Kenett et al., 2014). Semantic memory network methods have proven valuable in studying these processes (Benedek et al., 2017; Bernard et al., 2019; Bieth et al., 2021; Ovando-Tellez et al., 2022). Semantic networks consist of a set of nodes, which represent concepts, or words, interconnected by links that represent the strength of the semantic association between them. Semantic networks provide a structure on which (censored or biased) random walk approaches have been tested to mimic semantic memory search (Zemla & Austerweil, 2017). (Pseudo-)random walks on a semantic network mimic paths that can be taken into the network to move from one node to another one. The use of those models was essentially used to explaining fluency tasks (Abbott et al., 2015) and memory retrieval of remote associates (Kenett & Austerweil, 2016), but they have not yet been combined with decision models that could bring new insights into how individuals reach a creative solution. Based on this literature, we modeled the *explorer* module as a random walk wandering into semantic networks.

Second, valuation and selection processes are typically studied using decision models. Utility (economic term for subjective value) functions can well capture valuation of multi-attribute options that weigh attributes differently depending on individuals (Lopez-Persem et al., 2017; Samuelson, 1938; Von Winterfeldt & Fischer, 1975). Hence, we modeled the *valuator* module of our model as a utility function that assigns subjective values to candidate ideas based on the subjective evaluation of their adequacy and originality, considered the necessary attributes of a creative idea.

Third, the computed subjective value is then used to make a decision. Simple decision models like *softmax* functions (Luce, 1959) can explain many types of choices, ranging from concrete food choices to abstract moral choices, as soon as they rely on subjective values. Briefly, a softmax function is a mathematical function that convert a decision-value, i.e., the subjective values of options, into a probability of choosing one option or another. Here, we reasoned that such a simple function could capture and predict creative choices (*selector* module) when taking subjective values of candidate ideas as input.

Overall, through different approaches to test our hypotheses, we developed an original computational model (Figure 1) in which each module (*explorer*, *valuator*, *selector*) was modeled separately. We aimed at 1) determining whether subjective valuation occurs during idea generation (creativity task) and defining a *valuator* module from behavioral measures during the decision-making tasks; 2) Developing the *explorer* and *selector* modules, and characterizing which module(s) relies on subjective valuation (*explorer* and/or *selector*); 3) Simulating surrogate data from the full model composed of the three modules and comparing it to human behavior; and 4) assessing the relevance of the model parameters for creative abilities.

## Empirical study

### Methods

#### Participants

An official ethics committee approved the study (CPP Ouest II – Angers). Seventy-one participants were recruited and tested thanks to the PRISME platform of the Paris Brain Institute (ICM). They gave informed consent and were compensated for their participation. Inclusion criteria were: being right-handed, native French speakers, between 22 and 40 years old, with correct or corrected vision, and having no history of neurological or psychiatric disease. Two participants were excluded because of a misunderstanding of the instructions, bringing the final number of participants to 69 (41 females and 28 males; mean age: 25.8±4.5; mean level of education: number of study years following French A-levels: 5.0±1.6). The initial sample size was defined based on the interindividual correlations that we wanted to address between the model parameters and the creativity scores from the battery of tests. Using the software G*Power, we estimated that to detect a positive moderate effect size (r=0.3) with a statistical power of 80%, for a p-value threshold of 0.05, we needed 64 participants. As we anticipated outliers and potential exclusions, we planned to include 75 participants but four did not show up for their appointment, resulting in 71 included participants.

#### Experimental Design

Each participant performed three types of tasks of creative generation and evaluation of ideas, which were followed by a battery of tests classically used in the laboratory and assessing the participant’s creative abilities. All tasks and tests were computerized and administered in the same fixed order for all participants.

##### Free Generation of Associations Task (FGAT)

The Free Generation of Associations Task (hereafter referred to as FGAT) is a word association task, previously shown to capture aspects of creativity (Bendetowicz et al., 2017; Prabhakaran et al., 2014). It is composed of two conditions, presented successively, always in the same order. Cue words selection is detailed in SI.

##### FGAT-First Condition

After a 5-trials training session, participants performed 62 trials of the first condition block (hereafter referred to as FGAT-first). They were presented with a cue word and instructed to provide the first word that came to mind after reading it. They had 10 seconds to find a word and press the spacebar and then were allowed 10 seconds maximum to type it on a keyboard. This condition was used to explore the participants’ spontaneous semantic associations and served as a control condition that is not a creative task per se.

##### FGAT-Distant Condition

In a different following block, participants were administered 62 trials of the second condition of the task (hereafter referred to as FGAT-distant). On each trial, they were presented with a cue word as in the previous condition and instructed to press the spacebar once they had thought of a word unusually associated with the cue. They were asked to find a distant but understandable associate and to think creatively. They had 20 seconds to think of a word, press the spacebar, and then were allowed 10 seconds maximum to type it. This condition measures the participants’ ability to intentionally produce remote and creative associations.

##### Rating Tasks

After the FGAT task, participants performed two rating tasks. In the first block, they had to rate how much they liked an association of two words (likeability rating task). Then, in a separate block performed after the Choice task (see below), they had to rate the originality and the adequacy (originality and adequacy rating task) of the same associations as in the likeability rating task.

##### Likeability Rating Task

After a 5-trial training session, participants performed 197 trials in which they were presented with an association of two words (cue-response, see below) and asked to rate how much they liked this cue-response association in a creative context, i.e., how much they liked it or would have liked to find it during the FGAT *Distant* condition. A cue-response association was displayed on the screen, and 0.3 to 0.6 seconds later, a rating scale appeared underneath it. The rating scale’s low to high values were represented from left to right, without any numerical values but with 101 steps and a segment indicating the middle of the scale (later converted in ratings ranging between 0 and 100). Participants entered their rating by pressing the left and right arrows on the keyboard to move a slider across the rating scale, with the instruction to use the whole scale. Once satisfied with the slider’s location, they pressed the spacebar to validate their rating and went on to the subsequent trial. No time limit was applied, but participants were instructed to respond as spontaneously as possible. A symbol (a heart for likeability ratings) was placed underneath the scale as a reminder of the dimension on which the words were to be rated.

##### Originality and Adequacy Ratings

The originality and adequacy rating task was performed after the likeability rating task and the choice task to avoid any prior influence of these dimensions on the likeability ratings and choices. After a 5-trial training session, participants performed a block of 197 trials. They were asked to rate the same set of associations as in the likeability task, but this time in terms of originality and adequacy, and in a different random order. The instructions described an original association as ‘original, unusual, surprising’. An adequate association was described as ‘appropriate, understandable meaning, relevant, suitable’. Note that the instructions were given in French to the participants and the adjectives used here are the closest translation we could find.

For each cue-response association, participants had to rate originality and adequacy dimensions one after the other, in a balanced order (in half of the trials, participants were asked to rate the association’s adequacy before its originality, and in the other half of the trials, it was the opposite). The order was unpredictable for the participant. Similar to the likeability ratings, the rating scale appeared underneath the association after 0.3 to 0.6 seconds, with a different symbol below it: a star for originality ratings and a target for adequacy ratings, as depicted in Figure 2.

**Figure 2:**
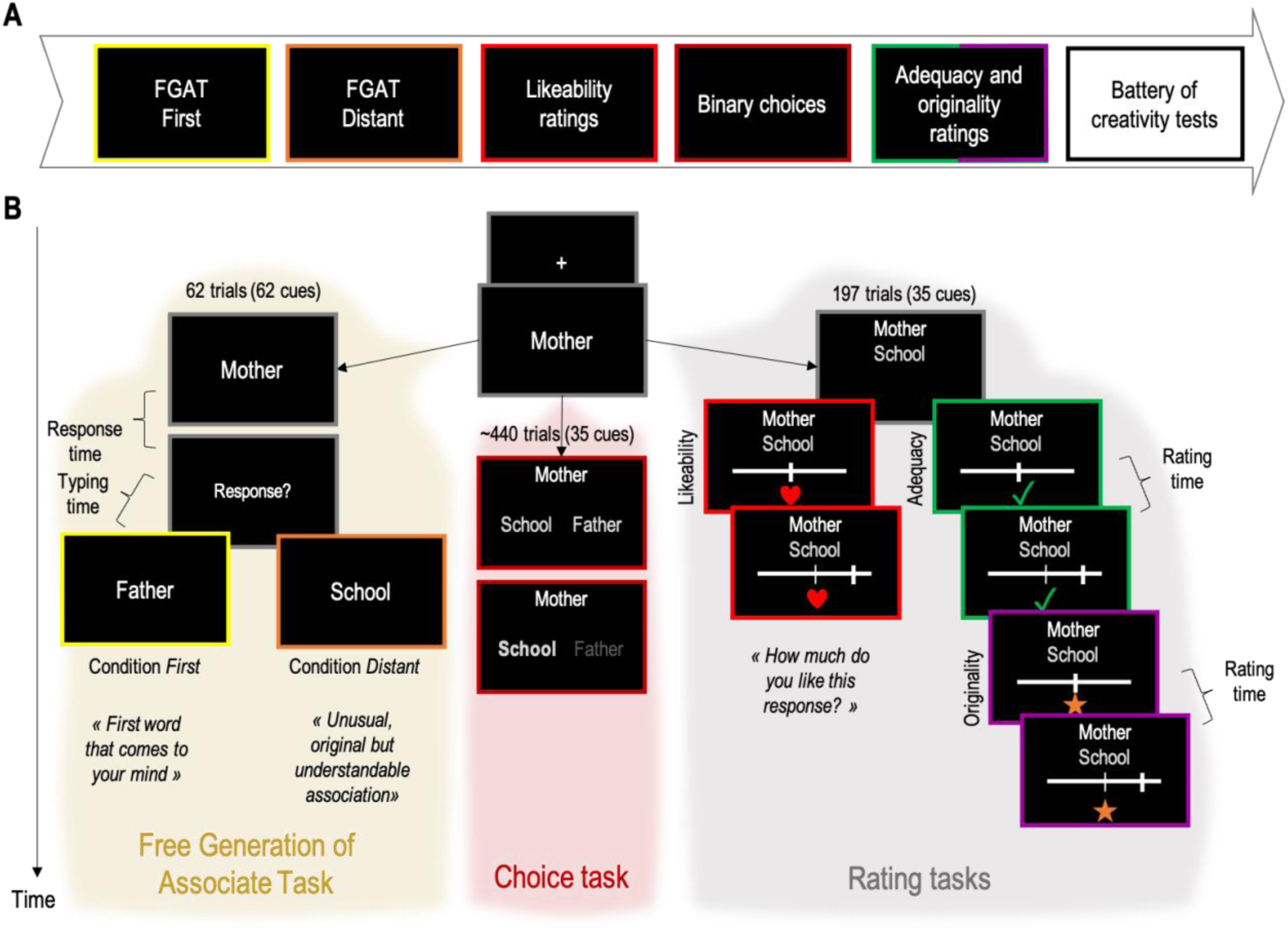
Experimental design. **A.** Chronological order of successive tasks. **B**. From top to bottom, successive screen shots of example trials are shown for the three types of tasks (left: FGAT task, middle: choice task, right: rating tasks). Every trial started with a fixation cross, followed by one cue word. In the **FGAT** task, when participant had a response in mind, they had to press the space bar and the word “Response?” popped out on the screen. The FGAT task had two conditions. Participants had to press a space for providing the first word that came to their mind in the *First* condition and an unusual, original but associated word in the *Distant* condition. In the **choice** task, two words were displayed on the screen below the cue. Participants had to choose the association they preferred using the arrow keys. As soon as a choice was made, another cue appeared on the screen and the next trial began. In the **rating** tasks, one word appeared on the screen below the cue. Then a scale appeared on the screen, noticing subjects that it was time for providing a response. In the likeability rating task, participants were asked to indicate how much they liked the association in the context of FGAT-distant. In the adequacy and originality rating tasks, each association was first rated on either adequacy and originality and then on the remaining dimension. Order was counterbalanced (see Methods for details).

##### Cue-Word Associations in the Rating Tasks

The 197 cue-response associations presented in the rating were built with 35 FGAT cue words randomly selected for each participant after they performed the FGAT task. We used a MatLab script that implemented an adaptive design with the following rules. Each of the 35 cue words was paired with seven different words, amounting to 245 possible associations in total. We paired each cue word with 1) the participant response to the cue FGAT First, 2) the participant’s response to the cue in FGAT Distant, 3) one word selected randomly from the most common FGAT first responses from another dataset collected previously in the lab that gathers the responses of 96 independent and healthy participants on a similar FGAT task, 4) one word selected randomly from the less common FGAT First responses from this other dataset, 5) one word selected randomly from the most common FGAT Distant responses from the same other dataset, 6) one word selected randomly from the less common FGAT Distant responses from the same other dataset, and 7) one unrelated association for each cue (‘cow’ with ‘inverse’ for instance) (See SI Supplementary Methods for a full description). We used these word associations from another study and unrelated associations to obtain a sufficient sampling of all possible combinations of adequacy and originality ratings (to estimate likeability with sufficient statistical power).

##### Choice Task

Participants performed a binary choice task between the likeability rating task and the adequacy-originality rating task. They had to choose between two words the one they preferred to be associated with a cue in a creative context, i.e., in the FGAT *Distant* context. Instructions were as follows: ‘For example, would you have preferred to answer “silver” or “jewelry” to “necklace” when generating original associations during the previous task?’ (There was additionally a reminder of the FGAT *Distant* condition, in the instructions). Details of the task and how the items were selected can be found in SI Supplementary Methods.

##### Battery of Creativity Tests

A battery of creativity tests and questionnaires run on Qualtrics followed the previous tasks to assess the participants’ creative abilities and behavior. It was composed of the alternative uses task (AUT), the inventory of Creative Activities and Achievements (ICAA), a self-report of creative abilities, a scale of preferences in creativity between adequacy and originality (SPC), and a fluency task on six FGAT cues. They are described in detail in the Supplementary Methods.

#### Statistical Analysis

All analyses were performed using Matlab (MATLAB. (2020). 9.9.0.1495850 (R2020b). Natick, Massachusetts: The MathWorks Inc.).

##### FGAT Responses

The main behavioral measures in the FGAT task are the response time (pressing the space key to provide an answer), the typing speed (number of letters per second), and the associative frequency of the responses. This frequency was computed based on a French database called *Dictaverf* (http://dictaverf.nsu.ru/)(Debrenne, 2011) built on spontaneous associations provided by at least 400 individuals in response to 1081 words (each person saw 100 random words). Frequencies were log-transformed to take into account their skewed distribution toward 0. Cues varied in terms of steepness (the ratio between the associative frequency of the first and second free associate of a given cue word), which was a variable of interest. Subjects’ ratings of their responses (adequacy, originality, and likeability) were also used as variables of interest.

Linear regressions were conducted at the subject level between normalized variables. Significance was tested at the group level using one sample, two-tailed t-tests on coefficient estimates.

##### Likeability Ratings Relationship with Adequacy and Originality Ratings

In this analysis, we aimed at explaining how likeability ratings integrated adequacy and originality dimensions. We tested whether this integration was linear or not (with exponential terms or with the addition of interaction terms, or without) and whether adequacy and originality were in competition or not (one relative weights balancing adequacy and originality or two independent weights).

First, we fitted 12 different functions to likeability ratings capturing different types of relationships (for instance linear of not linear between likeability (L) and adequacy (A), and originality (O):

- Linear models: 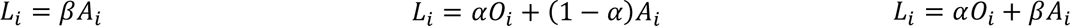
- Linear with interaction term models: 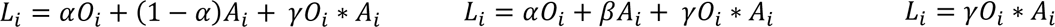
- Non-linear models (with the same non-linearity on both dimensions): 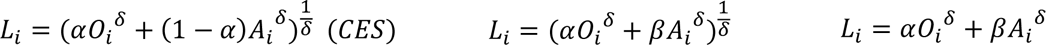
The first non-linear model is also referred to as Constant Elasticity of Substitution (CES) (Andreoni & Miller, 2003) - Non-linear models (with different non-linearity on both dimensions): 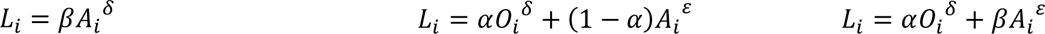

Greek letters correspond to free parameters estimated with the fitting procedure described below; i refers to a given cue-response association.

Then, we compared the performance of the 12 models to explain the relationship between likeability ratings and adequacy and originality ratings. Model fitting and comparison procedure is detailed in Methods *Model Fitting and Comparison*.

### Results

Sixty-nine subjects were included in the analyses (see Methods *Participants*). The experiment consisted of several successive tasks (Figure 2, see Methods *Experimental Design*): the Free Generation of Associate Task (FGAT), designed to investigate generative processes and creative abilities, a likeability rating task, a choice task, an originality, and adequacy rating task, and a battery of creativity assessment.

#### FGAT Behavior: Effect of Task Condition on Speed and Link with Likeability

In the *First* condition of the FGAT task, participants were asked to provide the first word that came to mind in response to a cue. In the *Distant* condition, they had to provide an original, unusual, but associated response to the same cues as in the *First* condition (see Figure 2 and Methods *Free Generation of Associations Task (FGAT)*).

We investigated the quality and speed of responses in the FGAT task in the *First* and *Distant* conditions. The quality of responses was investigated using their associative frequency obtained from the French database of word associations *Dictaverf* (see Methods *Statistical Analysis*), and using the ratings that participants provided in three rating tasks requiring them to judge how much they liked an idea (likeability of a response to the FGAT *Distant* condition, see Methods *Rating Tasks*), how much original they found it (originality), and how appropriate (adequacy).

##### FGAT Responses: Associative Frequency

Consistent with the instructions of the FGAT conditions, we found that participants provided more frequent responses (i.e., more common responses to a given cue based on the French norms of word associations *Dictaverf*) in the *First* condition than in the *Distant* condition (log(Frequency_First_)=-3.25±0.11, log(Frequency_Distant_)=-6.21±0.11, M±SEM, t(68)=18.93, p=8.10^-29^). Then, we observed that response time in the FGAT task decreased with the cue-response associative frequency, both in the *First* (β=-0.34±0.02, t(68)=-15.92, p=1.10^-24^) and *Distant* (β=-0.10±0.02, (68)=-6.27, p=3.10^-8^) conditions, suggesting that it takes more time to provide a rare response compared to a common one (Figure S1A). We also observed that the cue steepness (how strongly connected is the first associate of the cue, see Methods *Statistical Analysis*) also significantly shortened response time for *First* responses but not significantly for *Distant* responses (β_First_=-0.13±0.02, t(68)=-8.5, p=3.10^-12^; β_Distant_=-0.02±0.01, t(68)=-1.16, p=0.25, Figure S1B).

##### FGAT Responses: Adequacy and Originality

Using adequacy and originality ratings provided by the participants, we found that *First* responses were rated as more adequate than *Distant* responses (Adequacy_First_=86.47±0.99, Adequacy_Distant_=77.24±1.23, t(68)=9.29, p=1.10^-13^), but *Distant* responses were rated as more original than *First* responses (Originality_First_=33.80±1.74, Originality_Distant_=64.43±1.37, t(68)=-16.36, p=3.10^-25^). Note that the difference in originality ratings (*First* versus *Distant* responses) was greater than the difference in adequacy ratings (t(68)=-13.87, p=2.10^-21^), suggesting that *Distant* responses were found both adequate and original, i.e., creative, while *First* responses were mainly appropriate (Figure 3A).

**Figure 3:**
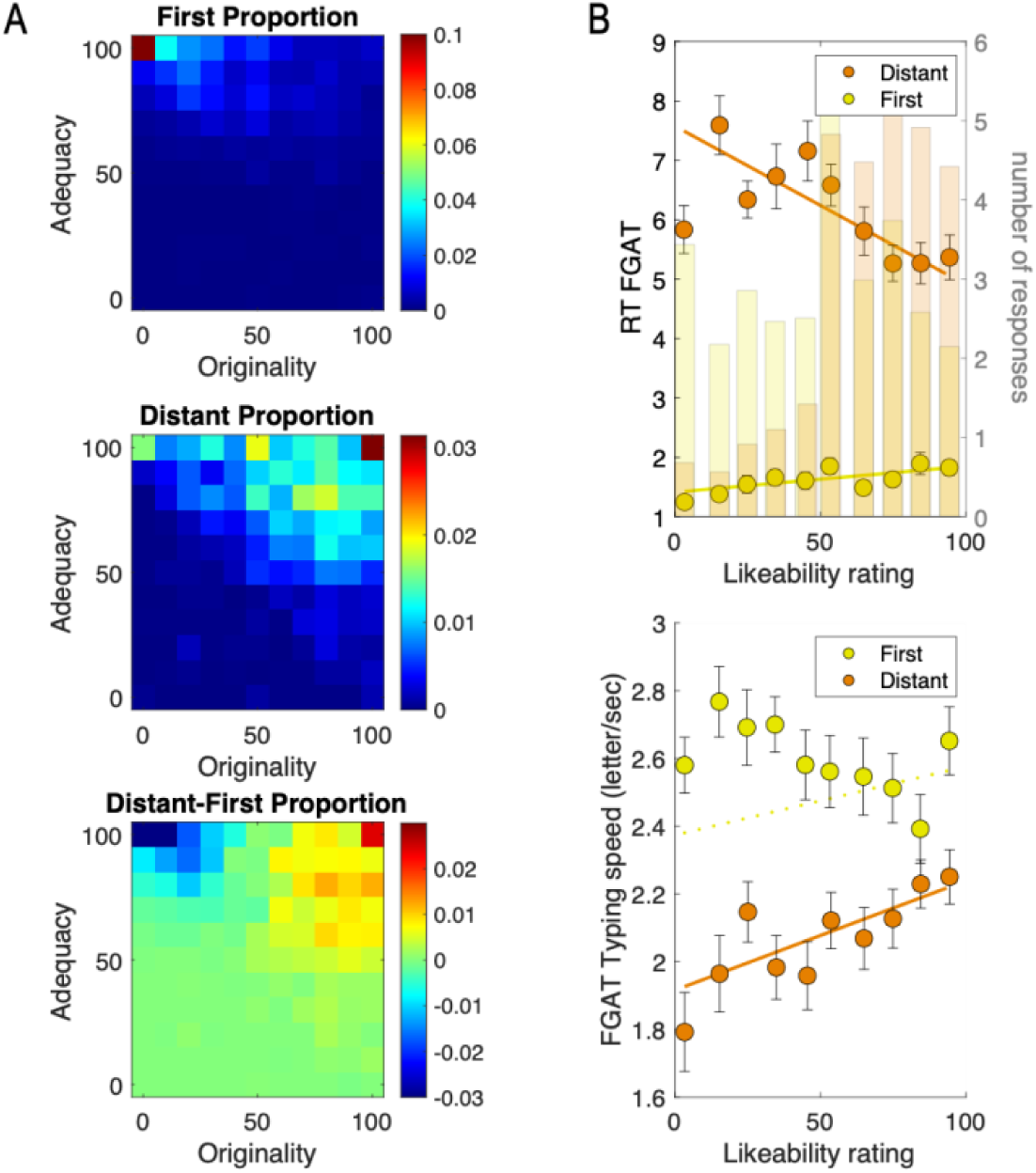
Behavioral results of the FGAT task. **A.** Heatmaps of First (top), Distant (middle) and Distant-First (bottom) proportions of responses per bin of adequacy and originality ratings. **B**. Correlation between response time (top) and typing speed (bottom) in the FGAT task and likeability ratings of the FGAT responses for the *First* (yellow) and *Distant* (orange) conditions. Circles indicate binned data averaged across participants. Error bars are intersubject s.e.m. Solid lines correspond to the averaged linear regression fit across participants, significant at the group level (p<0.05). Dotted lines indicate that the regression fit is non-significant at the group level (p>0.05). In **B top**, transparent bars correspond to the average number of responses per bin of likeability.

##### FGAT Responses: Likeability

Last, we considered that response time and typing speed could reflect an implicit valuation of responses (Niv, 2007). To test whether an implicit subjective valuation of responses happened during the FGAT creative condition (*Distant*), we investigated the link between response time, typing speed, and the likeability of their own FGAT responses (see Methods *Statistical Analysis*). We found that response time in the *Distant* condition decreased with likeability (β_Distant_=-0.15±0.02, t(68)=-7.25, p=5.10^-10^) and that typing speed increased with it (β_Distant_=0.08±0.02, t(68)=3.88, p=2.10^-4^). Participants were faster for providing *Distant* FGAT responses they liked the most. The pattern was different in the *First* condition, in which we observed a significant increase in response time with likeability (β_First_=0.08±0.02, t(68)=3.78, p=3.10^-4^) and no significant effect of likeability on typing speed (β_First_=0.009±0.02, t(68)=0.36, p=0.72). The effects of likeability significantly differed at the group level between the *First* and *Distant* conditions (*Distant* versus *First* effect of likeability on response time: t(68)=-7.30, p=4.10^-10^; on typing speed: t(68)=2.21, p=0.03, Figure 3B).

Note that the link between likeability rating and response time, or typing speed remains after removing confounding factors (adequacy and originality ratings, SI Table S1).

Together, those findings suggest that likeability might have been cognitively processed during the FGAT task and influenced the behavior, particularly during the FGAT *Distant* condition, which is assumed to require an evaluation of the response before the participants typed their answers. As a control analysis, we also found that likeability ratings drove choices (choice task, see SI Supplementary Results and Figure S2), suggesting that likeability is relevant both in the FGAT *Distant* condition, and in binary choices linked to creative response production. We next assessed how likeability ratings relied on adequacy and originality ratings.

#### Likeability Depends on Originality and Adequacy Ratings

To better understand how subjects built their subjective value and assigned a likeability rating to a cue-response association, we focused on the behavior measured during the rating tasks. In the rating tasks, participants judged a series of cue-response associations in terms of their likeability, adequacy and originality (see Figure 2 and Methods *Rating Tasks*). Here, we explored the relationship between those three types of ratings.

We first observed that likeability increased with both originality and adequacy (Figure 4). Then, to precisely capture how adequacy and originality contributed to likeability judgments, we compared 12 different linear and non-linear models (see Methods *Likeability Ratings Relationship with Adequacy and Originality Ratings*). Among them, the Constant Elasticity of Substitution (CES) model outperformed (Lopez-Persem et al., 2017) the alternatives (Estimated model frequency: Ef=0.36, Exceedance probability: Xp=0.87). CES combines originality and adequacy with a weighting parameter α and a convexity parameter δ into a subjective value (likeability rating) (see equation in Figure 1 and fit in Figure 4). In our group of participants, we found that α was significantly lower than 0.5, indicating an average overweighting of adequacy compared to originality (Mean α=0.43±0.03, t(68)=-2.37, p=0.02, one sample two-sided t-test against 0.5). Additionally, δ was significantly lower than 1, indicating that a balanced equilibrium between adequacy and originality was in average preferred compared to an unbalanced equilibrium, such as associations with high adequacy and low originality (Mean δ=0.62±0.11, t(68)=-3.46, p=9.10^-4^, one sample two-sided t-test against 1).

**Figure 4.**
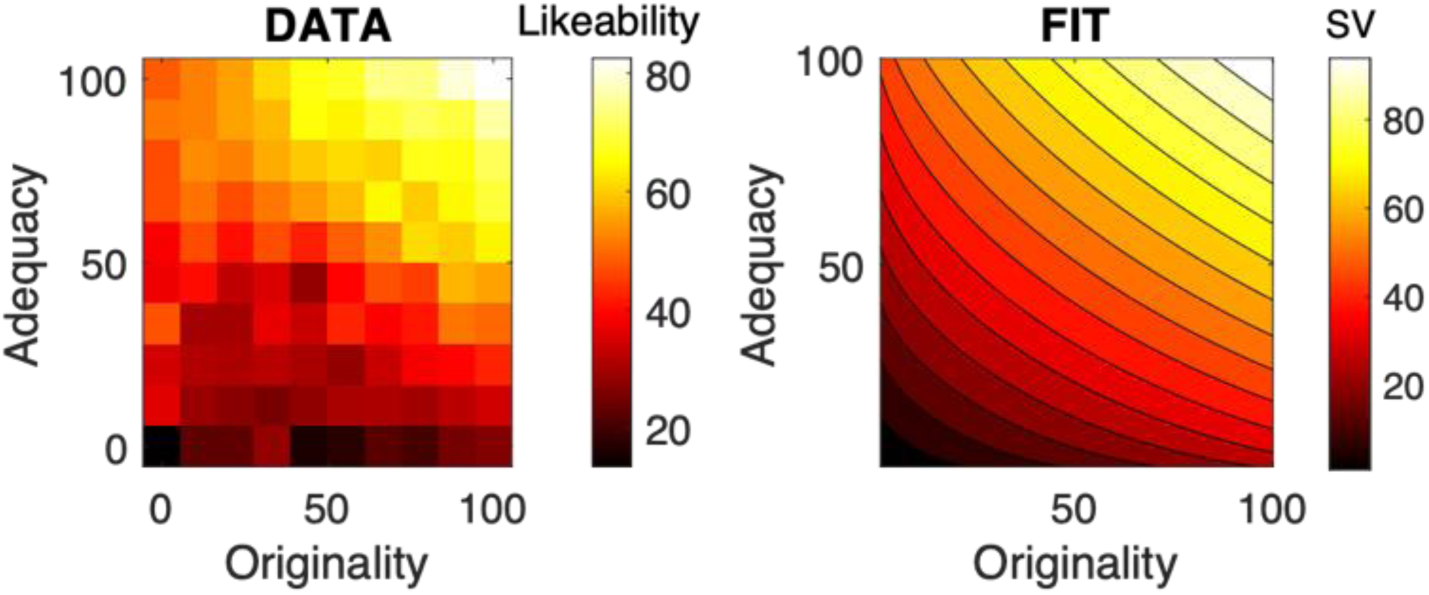
Behavioral results of the rating tasks: building the valuator module. Average likeability ratings (left) and fit (right) are shown as functions of adequacy and originality ratings. Black to hot colors indicate low to high values of likeability ratings (left) or fitted subjective value (SV, right). The value function used to fit the ratings was the CES utility function.

#### Individual Preferences and Responses Creativity

In the previous analyses, we found that the new ideas people like the most are produced the fastest. On the contrary, we found that infrequent ideas took more time to be provided. Unsurprisingly, when assessing the relationship between frequency of responses and likeability ratings of *Distant* responses in our group of participants, we found no significant effect at the group level (linear regression of likeability ratings against frequency at each individual level, one sample two-sided t-test at the group level on the mean regression coefficient: t(68)=0.13, p=0.89, Figure S3).

Nevertheless, in the previous analyses, we also found that preferences rely on a balance between adequacy and originality. We then checked the relationship between frequency and likeability of *Distant* responses by splitting our group of participants according to the value of the α parameter. Participants with α>0.5 (favoring originality in their likeability judgments) were pooled in Group 1 and participants with α <0.5 (favoring adequacy in their likeability judgement) in Group 2. We found that Group 1 preferred (rated likeability higher) more creative ideas (t(28)=-2.70, p=0.01, figure S3), while Group 2 preferred less creative ideas (t(39)=2.60, p=0.013, figure S3). The difference of regression coefficient between groups was strong and significant (two-samples, one-sided t-test: t(67)=4.23, p=7.10^-5^). In other word, the link between likeability and creativity was positive only in participants who favored originality over adequacy.

To go a step further, we tested whether ideas provided by Group 1 during FGAT *Distant* were overall less frequent than *Distant* ideas provided by Group 2. The comparison was significant (two-samples, one-sided t-test: t(67)=-1.812, p=0.037).

To summarize, individuals who favor originality in their likeability ratings prefer more creative ideas and provide more creative ideas, compared to individuals who favor adequacy.

### Discussion

The first aim of our study was to determining whether subjective valuation occurs during idea generation and defining a valuator module from the decision-making tasks. Overall, these results indicate that subjective valuation occurs during idea generation, as we observed significant relationships between response speed and likeability ratings in the generation task, with preferred responses being provided faster. This result can be interpreted as a form of behavioral energization, which mechanisms need to be better understood. The choice task allowed us to verify that likeability was the most relevant dimension that participants used to choose between options, consistent with previous studies on value-based decision-making (Lopez-Persem et al., 2017, 2020).

The rating tasks have allowed us to characterize how likeability is built from the adequacy and originality of ideas. Overall, participants overweighted adequacy (weight parameter) and preferred responses with balanced originality and adequacy compared to unbalanced responses (convexity parameter). This result is in line with previous literature showing that originality tends to be openly or theoretically valorized but depreciated in practice (Blair & Mumford, 2007; Mueller et al., 2012). Nevertheless, it is essential to highlight here that participants overall take into account both dimensions, but vary in the way they do it: some individuals favor high originality over high adequacy in their likeability judgment (high α parameter), while others favor equilibrium between the two dimensions (delta lower than 1). Importantly, we found that this equilibrium (through the α parameter) seems to be influential in participant’s creativity: participants overweighting originality in their preference provide less frequent ideas, and thus more creative ideas.

The utility function fitting also constitutes the development of the valuator module in our general computational model, as the Constant Elasticity of Substitution utility function (CES), that builds a subjective value from adequacy and originality ratings.

In the next section, we will address the other aims of this study and develop a computational model that aims at disentangling how valuation differentially impacts exploration and selection processes underlying creative ideation. Two non-exclusive alternative hypotheses exist. Valuation either influence the exploration phase: navigating from one idea to another when searching for a creative idea is biased by preferences, or the selection phase: among the considered ideas, the one with the highest likeability is selected.

## Computational Modeling of Empirical Data

### Methods

To develop our computational model, we focused on its three modules separately. The explorer module was developed using simulations with semantic networks, and the valuator and selector modules were developed using model fitting and model comparisons. Model simulations aims at generating surrogate data that are then analyzed and compared to human data. Model fitting aims at adjusting parameters of equations at the individual level to match the data. Model comparison aims at determining which equation better matches the data (at the group level), once the parameters have been estimated.

We first explain below the model fitting and comparison procedures that we used. Then, we explain how we modelled the valuator (partially based on analyses conducted in the empirical study) for all participants.

Then, as the second aim of the current study was to identify whether likeability influences exploration or selection, and to develop the full model, we explain how we simulated data from various versions of the explorer, and how we developed the selection module. Next, to address the third aim of this study, we combined the three modules to get a ‘full’ model and generated surrogate data to compare the model behavior to participants’ behavior.

Finally, to assess the relevance of model parameters to creative abilities (fourth aim), we conducted a canonical correlation analysis.

#### General Procedure for Model Fitting and Comparison

Every model/module was fitted at the individual level to ratings and choices using the Matlab VBA-toolbox (https://mbb-team.github.io/VBA-toolbox/), which implements Variational Bayesian analysis under the Laplace approximation (Daunizeau et al., 2009; Stephan et al., 2009). This iterative algorithm provides a free-energy approximation to the marginal likelihood or model evidence, which represents a natural trade-off between model accuracy (goodness of fit) and complexity (degrees of freedom) (Friston et al., 2007; Penny, 2012). Additionally, the algorithm provides an estimate of the posterior density over the model free parameters, starting with Gaussian priors. Individual log-model evidence were then taken to group-level random-effect Bayesian model selection (RFX-BMS) procedure (Rigoux et al., 2014; Stephan et al., 2009). RFX-BMS provides an exceedance probability (Xp) that measures how likely it is that a given model (or family of models) is more frequently implemented, relative to all the others considered in the model space, in the population from which participants were drawn (Rigoux et al., 2014; Stephan et al., 2009).

We conducted the first model comparison to determine which variable (Adequacy A, Originality O or Likeability L) best explained choices (SI Methods *Relationship Between Choices and Ratings*). The second model comparison was performed to identify which utility function (*valuator* module) best explained how originality and adequacy were combined to compute likeability (Methods *Likeability Ratings Relationship with Adequacy and Originality Ratings*). The third one aimed at establishing relationships between adequacy and originality ratings and associative frequency of cue and responses (Methods *Valuator Module: Combining Likeability, Originality, and Adequacy of the Rating Tasks with Responses Associative Frequency).* The fourth one aimed at identifying the best possible input variable for the *selector* module (Methods *Decision Functions as the Selector Module*).

#### Valuator Module: Combining Likeability, Originality, and Adequacy of the Rating Tasks with Responses Associative Frequency

For all participants, the ratings were used to estimate the likeability of a given response to a cue from its adequacy and originality (Methods *Likeability Ratings Relationship with Adequacy and Originality Ratings)*, themselves estimated from its associative frequency.

We investigated how adequacy and originality were linked to associative frequency between a cue and a response F_ci_. We tested for linear and non-linear relationships between adequacy/originality and frequency using polynomial fits of second order. For each dimension X (A or O), we compared three models:

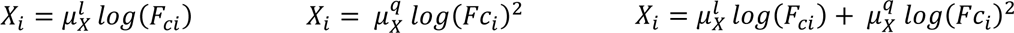

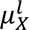 corresponds to the linear regression coefficient and 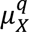 to the quadratic regression coefficient.

#### Model Identification Group and Test Group

For the next analyses, we randomly split our group of participants into two subgroups, one group to develop the *explorer* and *selector* modules (2/3 of the group: 46 subjects) and one group to validate the full model (combination of the *explorer*, *valuator* and *selector* modules) by comparing its behavioral prediction to the actual behavior of the participants (23 subjects).

#### Modeling the Explorer Module

We modeled the explorer module following a three-step procedure. First, we built semantic networks (for each cue) from a database available online to which we added the participant’s responses. Then, we developed random walks that would wander into those networks according to different rules (biased by associative frequency or likeability, for instance).

Finally, we compared the probabilities of those random walks to reach the *First* and *Distant* responses (nodes) of each participant for each cue during their trajectories in the semantic networks.

##### Construction of Semantic Networks

For each FGAT cue, we built a semantic network based on the Dictaverf database and the FGAT responses from the current dataset. Each network corresponds to an unweighted and undirected graph (an edge linked two nodes if the frequency of association between them was higher than 0). See details in SI Methods.

##### Random Walks Variants and Implementation

We used censored random walks that start at a given cue and walk within their associated network N. Censored random walks have the property of preventing return to previously visited nodes. In case of a dead-end, the censored random walk starts over from the cue but does not go back to previously visited nodes. The five following variants of censored random walks were applied to the semantic networks to simulate potential paths.

- The random walk random (RWR) was a censored random walk starting at the cue and with uniform distribution of probabilities of transition from the current node to each of its neighbors (excluding previously visited nodes).
- The random walk frequency (RWF) was a censored random walk biased by the associative frequency between nodes, where the probability of transition from one node to another one is defined as follows:

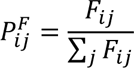

with 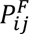 the probability of transition to node j, 𝐹_𝑖𝑗_ the frequency of the association (in the C matrices described in the Supplementary Methods) with the current node i, and j all the other nodes linked to the current node n.
- Three additional censored random walks were run. They were biased by adequacy (RWA), originality (RWO), or likeability (RWL) of association between nodes and cue, where the probability of transition from one node to another one is defined as follows:

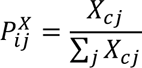

with 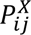 the probability of transition from node i to node j, 𝑋_𝑐𝑗_ the estimated adequacy, originality, or likeability of the node j with the cue node c, j are all the nodes linked to the current node i.

Estimated adequacy, originality, and likeability of all the network nodes (𝑋_𝑖_) were computed based on the model comparison results performed in the first section (see Methods *Valuator*

Module: Combining Likeability, Originality, and Adequacy of the Rating Tasks with Responses Associative Frequency). The following equations were consequently used:

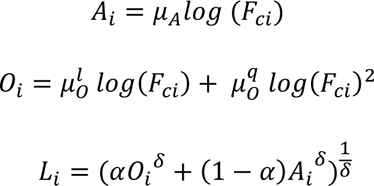

With 𝐹_𝑐𝑖_ as the frequency of association between the node and the cue (and see Supplementary Methods).

The number of steps performed by each random walk was constant across cues and participants and was defined by the median fluency score among the group, i.e., 18 steps, resulting in no more than 17 visited nodes.

##### Probability of Reaching First and Distant Responses for Each Participant and Cue

We computed the probability of reaching the *First* and *Distant* responses (Targets T) from a starting node cue (c) for each type of random walk as follows:

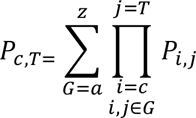

With G representing all possible paths between c and T, ranging from the shortest one (a) to the longest one (z) (limited to 18 steps) and i and j all pairs of nodes belonging to each path, linked by a transition probability P_i,j_. In other words, it corresponds to the sum of the cumulative product of edge weights for all the possible paths between the cue and the target shorter than 18 steps.

#### Decision Functions as the Selector Module

Next, we intended to decipher the criteria determining the selection of a given response. We compared seven criteria: random values, node rank (first visited nodes have higher chances of being selected), estimated adequacy, estimated originality, interaction between estimated adequacy and originality, sum of estimated adequacy and originality, and estimated likeability.

For each subject and cue, we simulated RWF as described above and retained the paths that contained both the *First* and *Distant* response of the subject for further analyses (the number of excluded cues ranged between 0 and 31 trials over 62, M=9.04 trials, exclusion mainly due to missing responses from participants either in the FGAT *First* or *Distant* condition).

Using the VBA toolbox, we fitted the choices (Response *First* or Response *Distant*) that subjects made among the hypothetically visited nodes (obtained by RWF simulations) for each trial using the following *softmax* functions:

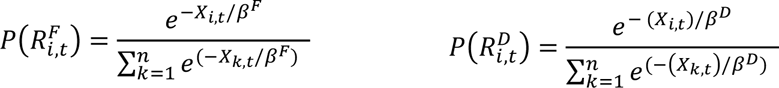

P is the probability of node *i* being selected as a response (R) in the *First* (F) or *Distant* (D) conditions for a given cue, among all the possible nodes *k* belonging to the *n* options from the paths at trial t. X corresponds to the values within seven different possible inputs (criteria defined earlier). 𝛽^𝐹^and 𝛽^𝐷^are free parameters estimated per subject, corresponding to the temperature (choice stochasticity).

We then compared the seven models for the *First* and *Distant* response separately and reported the results of the model comparison in the results. Details of the input structure is in Supplementary Methods.

#### Cross-Validation of the Model: Comparing the Surrogate Data to Human Behavior

To simulate the behavior of the remaining 23 subjects, we combined all the previously described modules together.

Concretely, we applied RWF with 18 steps on the built networks (see Methods *Construction of Semantic Networks*) and assigned values to each visited node according to each subject’s valuator module parameters. The list of visited nodes (candidate responses) for each cue and each subject was simulated without the constraint of containing participants’ *First* and *Distant* responses. The selection was made using an *argmax* rule on adequacy (winning criteria for the selector module) for the *First* response and on likeability (winning criteria for the selector module) for the *Distant* response (as we do not have the selection temperature parameters 𝛽^𝐹^and 𝛽^𝐷^ for those remaining subjects). We ran 100 simulations per individual following that procedure.

The rank in the path was used as a proxy for response time, and we analyzed surrogate data in the exact same way as subjects’ behavior.

For statistical assessment, regression estimates of ranks against frequency, steepness, and estimated likeability were averaged across 100 simulations per individual, and significance was addressed at the group level (one representative simulation was used in Figures 5 and S6). For this analysis, the group frequency of response was computed instead of *Dictaverf* associative frequency to 1) avoid any confounds with the structure of the graph, built with *Dictaverf*, and 2) compare the distribution of frequencies relative to the group.

**Figure 5.**
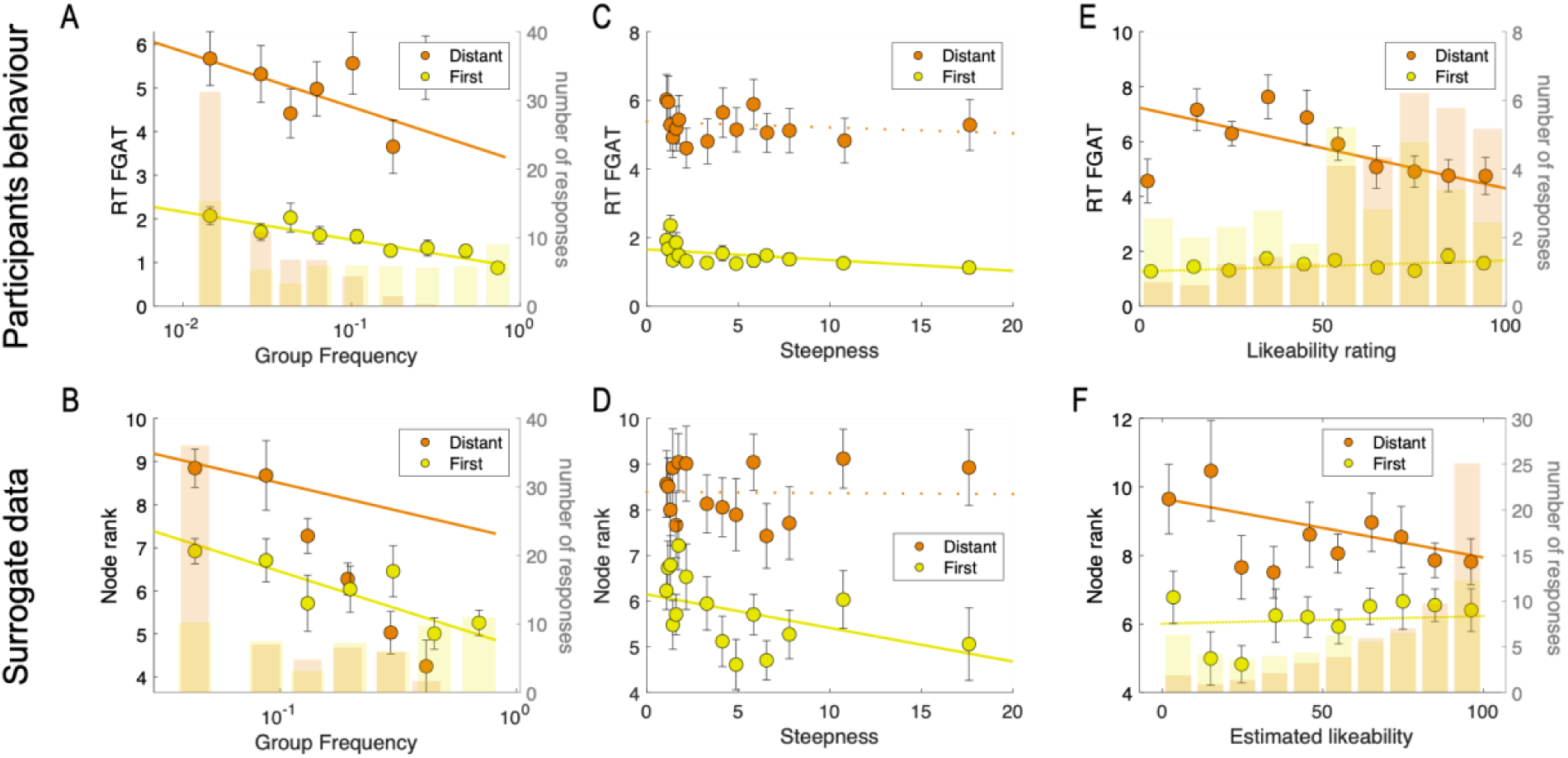
Response speed for the participants and surrogate data of the test group (n=23) **A, B.** Correlation between response time RT (A) or node rank (B) in the FGAT task and the response frequency for the *First* (yellow) and *Distant* (orange) conditions. **C, D.** Correlation between response time RT (C) or node rank (D) in the FGAT task and the cue steepness for the *First* (yellow) and *Distant* (orange) conditions. **E, F**. Correlation between response time RT (E) or node rank (F) in the FGAT task and likeability ratings (E) or estimated likeability (F) of the FGAT responses for the *First* (yellow) and *Distant* (orange) conditions. Circles indicate binned data averaged across participants. Error bars are intersubject s.e.m. Solid lines corresponds to the averaged linear regression fit across participants, significant at the group level (p<0.05). Dotted lines indicate that the regression fit is non-significant at the group level (p>0.05). In **A, B, E** and **F**, transparent bars correspond to the average number of responses per bin of frequency (A, B) or likeability (E, D). Note that the surrogate data presented in the Figure correspond to one simulation (among 100) that is representative of the statistics obtained over all simulations and reported in the text.

**Figure 6:**
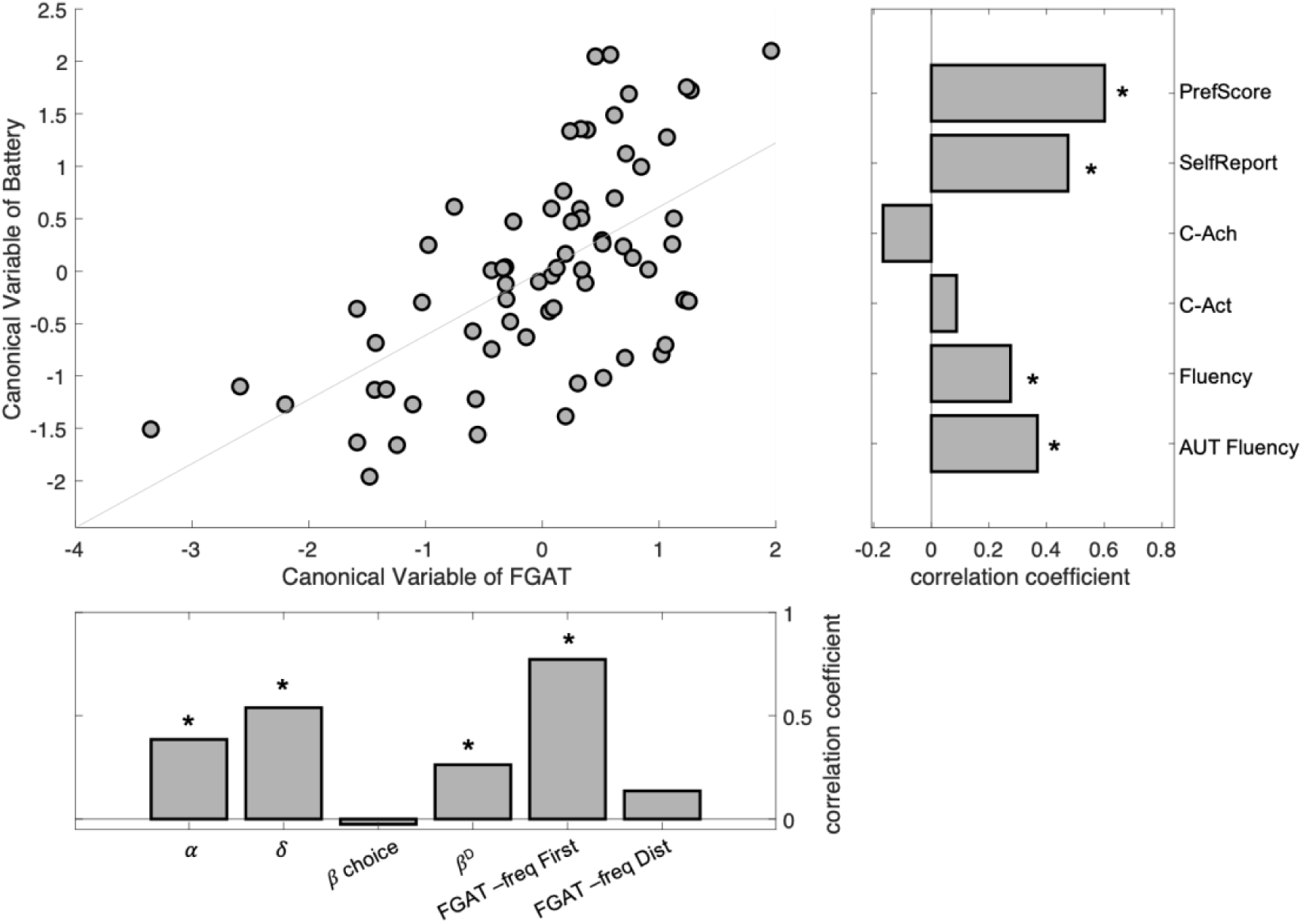
Canonical correlation between the FGAT parameters/metrics and creativity tests belonging to a battery. Top left. Correlation between the first canonical variables of the battery of tests and of the FGAT parameters/metrics. Each dot represents one participant. Top right: correlation coefficient between each battery test and the canonical variable of Battery. Bottom left: correlation coefficient between each FGAT parameters/metrics and the canonical variable of FGAT. Stars indicate significance (p>0.05).

#### Canonical Correlation Between Creativity Scores, FGAT Task, and Model Parameters

To investigate the link between creative abilities and our task and model parameters, we extracted the individual task scores and model parameters and grouped them into the label “FGAT scores and parameters”. We pooled the scores obtained from the battery of creativity test and labeled them “Battery scores”. We conducted a canonical correlation between those two sets of variables and checked for significance of correlation between the computed canonical variables of each set. Note that a canonical correlation analysis can be compared to a Principal Component Analysis, in the sense that common variance between two data sets is extracted into canonical variables (equivalent to principal components). Canonical variables extracted for each data set are ordered in terms of strength of correlations between the two data sets. Each variable within a data set has a loading coefficient that indicates its contribution to the canonical variable. Here, we extracted the coefficients of each variable on its respective canonical variable and reported them.

### Results

#### Computational Modelling of The Valuator Module

The goal of our computational model is to explain and predict the behavior of participants in the FGAT, by modeling an *explorer* that generates a set of candidate ideas, a *valuator* that assigns a subjective value to each candidate idea, and a *selector* that selects a response based (or not) on this subjective value. Our computational model thus needed to be able to predict the likeability of any potential cue-response associations, including those that have not been rated by our participants (see section *Valuator Module: Combining Likeability, Originality, and Adequacy of the Rating Tasks with Responses Associative* Frequency), and those that have not been expressed by participants during the FGAT *Distant* condition (hidden candidate ideas).

We found that adequacy and originality ratings could be correctly predicted by associative frequency (see SI Supplementary Results and Figure S4). Adequacy ratings could be well fitted through a linear relation with frequency (Ef_lin_=0.86, Xp_lin_=1), and originality could be estimated through a mixture of linear and quadratic links with frequency. This result allows us to estimate the adequacy and originality of any cue-response association for a given participant.

Importantly, we explored the validity of the *valuator* module using estimated adequacy and originality. We estimated likeability from the estimated adequacy and originality, using the individual parameters of the CES function mentioned above. We found a strong relationship between estimated and real likeability judgments (mean r=0.24±0.02, t(68)=11.04 p=8.10^-17^).

This result is not only a critical validation of our model linking likeability, originality, and adequacy, but also allows defining a set of parameters for each individual for the *valuator* module. Thanks to that set of parameters, we could significantly predict any cue-response association’s originality, adequacy, and likeability ratings based on its objective associative frequency. Henceforth, in the subsequent analyses, likeability, adequacy, and originality estimated through that procedure will be referred to as the “estimated” variables.

In the next section, using computational modeling, we address the second aim of our study, which was to develop the *explorer* and the *selector* and determine which module the *valuator* drives the most.

#### Computational Modelling of The Exploration and Selection Modules

##### Model Description and Overall Strategy

As we do not have direct access to the candidate ideas that participants explored before selecting and producing their response to each cue during the FGAT task, we adopted a computational approach that uses random walk simulations ran on semantic networks (one per FGAT cue) to develop the *explorer* module. We built a model that coupled random walk simulations (*explorer*) to a valuation (*valuator*) and selection (*selector*) function (Figure 1). The model takes as input an FGAT cue and generates responses for the *First* and *Distant* conditions, allowing us to ultimately test how similar the predicted responses from the model were to the real responses of the participants.

In the following analyses, we decompose the model into modules (random walks and selection functions) and investigate by which variable (estimated likeability, estimated originality, estimated adequacy, associative frequency, or mixtures) each module is more likely to be driven.

To assess the model’s validity, we developed it, conducted the analyses on 46 subjects (2/3 of them), and then cross-validated the behavioral predictions on the 23 remaining participants.

##### Modeling the Explorer Module Using Random Walks on Semantic Networks

For each cue, we built a semantic network from the *Dictaverf* database that was enriched from both *First* and *Distant* FGAT responses from all participants (see Methods *Construction of Semantic Networks*). Then, to investigate whether exploration could be driven by likeability, we compared five censored random walks (RW), each with different transition probabilities between nodes (random, associative frequency, adequacy, originality, or likeability, see Methods *Random Walks Variants and Implementation*). For each random walk, subject, and cue, we computed the random walk’s probability of visiting the *First* and the *Distant* responses nodes (Figure S5A). We found that the frequency-driven random walk (RWF) had the highest chance of walking through the *First* (mean probability = 0.30 ± 0.01; all p<10^-33^) and *Distant* (mean probability = 0.05 ± 0.004; all p<10^-4^) responses. This result suggests that the *explorer* module may be driven by associative frequency between words in semantic memory. According to this result, we pursued the analyses and simulations with the RWF as an *explorer* module for both *First* and *Distant* responses.

##### Visited Nodes with the RWF as a Proxy for Candidate Responses

To define sets of candidate responses that will then be considered as options by the *selector* module, we simulated the RWF model for each subject and each cue over 18 steps (see Methods *Probability of Reaching First and Distant Responses for Each Participant and Cue*). Each random walk produced a path: i.e., a list of words (nodes) visited at each iteration. Each node is associated with a rank (position in the path), which will then be used as a proxy of response time. As a sanity check, we compared the list of words obtained from those random walks to the participants’ responses to a fluency task on six FGAT cues (see Methods *Battery of Creativity Tests*). We identified the common words between the model path and the fluency responses for each subject. Then, using a mixed-effect linear regression with participants and cues as random factors (applied to both intercept and slope), we regressed the node model rank against its corresponding fluency rank. We found a significant fixed effect of the fluency rank (β= 0.12±0.03, t(649)=3.35, p=8.10^-3^, SI Figure S6), suggesting that those simulations provide an adequate proxy for semantic memory exploration.

Together, results reported in the two last sections suggest that a censored random walk driven by the frequency of word associations provides a good approximation of semantic exploration during response generation in the FGAT task and that likeability has a negligible role during that phase. Hence, valuation does not seem to play a significant role in the *explorer* module.

##### Modeling the Selector Module as a Decision Function

We then explored the possible factors driving individual decisions to choose a given response (*selector* module) among the word nodes visited by the *explorer* module. To investigate the selection of *First* and *Distant* responses among all nodes in each path, i.e., on which dimension responses were likely to be selected, we compared seven choice models with different variables as input: random values, node rank (first visited nodes have higher chances of being selected), estimated adequacy, estimated originality, interaction between estimated adequacy and originality, sum of estimated adequacy and originality, and estimated likeability (see Methods *Decision Functions as the Selector Module*). We found that estimated adequacy was the best criterion to explain the selection of *First* responses (Ef_adequacy_=0.89, Xp_adequacy_=1) and likeability was the best criterion to explain the selection of *Distant* responses (Ef_likeability_=0.66, Xp_likeability_=0.99) (Figure S5B). These results indicate that valuation (based on individual likeability) is needed to select a creative response in the creative condition of the FGAT (*Distant*).

### Validity of the Full Model: Does it Predict Behavioral Responses in the Test Group?

The next analyses address our third aim, to confront simulated and observed data. After having characterized the equations and individual parameters of the *valuator* on all participants using the rating tasks, and of the *explorer* and *selector* modules on a subset of participants (n_1_=46), we checked whether this model could generate surrogate data similar to the behavior of the remaining participants (test group, n_2_=23). We simulated behavioral data and response time from the full model (*explorer*, *valuator*, *selector*), depicted in Figure 1 (See Methods *Cross-Validation of the Model: Comparing the Surrogate Data to Human Behavior*).

We analyzed the behavior of the simulated data the same way we analyzed the behavior of the real human data of the test group. We found the same patterns at the group level (SI Table S2, Figure 5 and S6): 1) *First* responses were much more common than *Distant* responses (Figure 5A, B); 2) the rank in path decreased with the group frequency of responses, both for *First* and *Distant* responses (Figure 5A, B), confirming that it takes more time to provide a rare response compared to a common one; 3) Ranks decreased with the cue steepness, both for *First* and *Distant* responses (Figure 5C, D); 4) Ranks of the *Distant* responses decreased with estimated likeability. The effect was significant only for *Distant* responses, and the difference between regression estimates for *First* and *Distant* responses was significant. (Figure 5E, F); 5) *First* responses were more appropriate than *Distant* responses, but *Distant* responses were more original than *First* responses. The difference in originality rating between the *First* and *Distant* responses was bigger than the difference in adequacy (SI Figure S7).

Additionally, we checked whether the surrogate data generated by the model for each participant was relevant at the inter-individual level. We estimated the *selector* parameters for the test group and conducted the analyses on all participants to increase statistical power. We found that the mean response time per participant across trials of the FGAT *Distant* condition was correlated with the mean rank of *Distant* responses across trials in the model exploration path (r=0.72, p=1.10^-4^). Similarly, the mean associative frequency (*Dictaverf*) of participants’ *Distant* responses was significantly correlated with the mean frequency of the model *Distant* responses (r=0.53, p=9.10^-3^). These results mean that the model successfully predicted individual behavioral differences in the FGAT task.

### Relevance of Model Parameters for Creative Abilities

Finally, to address our fourth aim and assess the relevance of the individual model parameters in relation to the FGAT task for creative abilities, we defined two sets of variables: FGAT parameters and scores reflecting the *valuator*, *selector,* and *explorer* individual characteristics, and Battery scores related to several aspects of creativity (see Methods *Battery of Creativity Tests* and SI Methods). We conducted a canonical correlation analysis between those two sets in all participants and found one canonical variable showing significant dependence between them (r=0.61, p=0.0057). When assessing which variables within each set had the highest coefficient to the canonical score, we found that the two likeability parameters (α and δ, from the *valuator*), the inverse temperature (choice stochasticity, from the choice task, of the *Distant* response selection (from the *selector*) (see SI results and SI Methods *Relationship Between Choices and Ratings*) and the *First* response associative frequencies were significantly contributing the FGAT canonical variable. Additionally, fluency score from the fluency task and from the alternative uses task (AUT), creativity self-report, and PrefScore (self-report of preferences regarding ideas) significantly contributed to the Battery canonical variable. No significant contribution was observed from creative activities (C-Act) and achievements (C-Ach) in real-life scores (Table S3, Figure 6). Overall, this significant canonical correlation indicates that measures of valuation and selection relate to creative behavior.

### Discussion

Thanks to the computational modelling of empirical data, we addressed the second, third and fourth aims of our study, which were (2) developing the *explorer* and *selector* modules, and characterizing which module(s) relies on subjective valuation (*explorer* and/or *selector*); (3) simulating surrogate data from the full model composed of the three modules and comparing it to human behavior; and (4) assessing the relevance of the model parameters for creative abilities.

We have developed a computational model that includes three modules: an explorer, a valuator, and a selector. Through successive Bayesian model comparisons, we have found that the explorer is more likely to be driven by associative frequency of ideas than likeability of ideas, that the valuator integrates both adequacy and originality of ideas, and that the selector uses likeability to generate a final output to the creative idea generation. The model makes behavioral predictions that are accurate both at the group level (general relationship between response time and frequency of responses for instance), and at the individual level (given a set of valuation parameters specific to an individual, it predicts whether this individual will be fast or slow to provide creative responses for instance). Finally, the model parameters, together with the behavior in the FGAT, are predictive of creative abilities evaluated with a battery of creativity tests, suggesting that this model is relevant to creative abilities.

## General Discussion

Using data from an empirical study combining creativity tasks and decision-making tasks, as well as computational modelling from those data, we provided empirical and computational evidence in favor the involvement of subjective valuation in creativity. We found that subjective value energizes the participants behavior during idea generation, and is driving the selection of ideas (more than the exploration of ideas).

### Preferred Associations are Produced Faster when Thinking Creatively

Using the FGAT task, previously associated with creative abilities (Bendetowicz et al., 2017), we found that *Distant* responses were overall more original and slower in response time than *First* responses. In addition, response time decreased with steepness (only for *First*) and cue-response associative frequency. Those results are in line with the notion that it takes time to provide an original and rare response (Christensen et al., 1957; Beaty & Silvia, 2012).

Critically, we identified that the likeability of *Distant* responses was negatively linked to response time and positively linked to typing speed. Interpretation of response time can be challenging as it could reflect the easiness of choice (Ratcliff & Rouder, 1998), the quantity of effort or control required for action (Botvinick et al., 2001), motivation (Niv, 2007), or confidence (Ratcliff & Starns, 2009). In any case, this result, surviving correction for potential confounding factors (see Results *FGAT Behavior: Effect of Task Condition on Speed and Link with Likeability*), represents evidence that subjective valuation of ideas occurs during a creative (hidden) choice. To our knowledge, this is the first time that such a result has been demonstrated. With our computational model, we attempt to provide an explanation of a potential underlying mechanism involving value-based idea selection.

### Subjective Valuation of Ideas Drives the Selection of a Creative Response

The striking novelty our results reveal is the role of the *valuator* module coupled with the *selector* module in idea generation. These modules are directly inspired by the value-based decision-making field of research (Levy & Glimcher, 2012; Lopez-Persem et al., 2020). To make any kind of goal-directed choice, an agent needs to assign a subjective value to items or options at stake, so that they can be compared and one can be selected (Rangel et al., 2008). Here, we hypothesized that providing a creative response involves such a goal-directed choice that would logically require the subjective valuation of candidate ideas. After finding a behavioral signature of subjective valuation in response time and typing speed, we have shown that likeability judgments best explained *Distant* response selection among a set of options. This pattern was similar to the behavior observed in the choice task, explicitly asking participants to choose the response they would have preferred to give in the FGAT *Distant* condition. Assessing valuation processes during creative thinking is highly relevant to understanding the role of motivation in creativity, as decision-making research shows that valuation is closely related to motivation process, and it is assumed that subjective values energize behaviors (Pessiglione et al., 2007). Previous studies have highlighted the importance of motivation in creativity (Collins & Amabile, 1999; Fischer et al., 2019).

However, those reports were mainly based on interindividual correlations, while our study brings new evidence for the role of motivation in creativity with a mechanistic approach. Our model adds to this literature by demonstrating novel, precise, and measurable mechanisms by which motivation may relate to creative thinking at the intra-individual level. Through computational modeling of the empirical data, we showed how subjective valuation drove idea selection. We did not find that subjective valuation drove exploration better than associative frequency. This negative result does not exclude a potential role of motivation on the exploration phase of idea generation. Future investigations using for example individual semantic networks will be invaluable to confirm or deny the role of motivation and value-based decision-making in the exploration phase, as suggested by other authors (Lin & Vartanian, 2018). In any case, our findings support the hypothesis that the BVS (sometimes called the reward system) is involved in creative thinking and paves the way to later investigate its neural response during experimental creativity tasks.

Our study reveals some mechanisms about how individual preferences are built and used to make creative choices. We identified how originality and adequacy ratings were taken into account to build likeability, and determined preference parameters (relative weight of originality and adequacy and convexity of preference) to predict the subjective likeability of any cue-response association. Subjective likeability relies on subjective adequacy and originality. The identified valuation function linking likeability with adequacy and originality, i.e., the Constant Elasticity of Substitution utility function, has been previously used to explain moral choices or economic choices (Armington, 1969; Andreoni & Miller, 2003; Lopez-Persem et al., 2017), making it an appropriate candidate for the *valuator* module of our model. Overall, these results indicate that likeability is a relevant measure of the individual values that participants attributed to their ideas, and inform us on how it relies on the combination of originality and adequacy.

The second novelty of our study is to provide a valid full computational model composed of an *explorer*, a *valuator* and a *selector* module. We characterized these modules, and brought an unprecedented mechanistic understanding of creative idea generation. Moreover, this full model can generate surrogate data similar to real human behavior at the group and inter-individual levels.

### A Computational Model that Provides a Mechanistic Explanation of Idea Generation

The computational model presented in the current study is consistent with previous theoretical frameworks involving two phases in creativity: exploration and evaluation/selection (Campbell, 1960; Kleinmintz et al., 2019; Lin & Vartanian, 2018; Mekern et al., 2019; Simonton, 1998; Sowden et al., 2015). The *explorer* module was developed using random walks as it had been successfully done in previous studies to mimic semantic exploration (Austerweil et al., 2012; Kenett & Austerweil, 2016). Here, we found that the simulated semantic exploration was driven by associative frequency between words, but was not biased by subjective judgments of likeability, adequacy or originality. This result is consistent with the associative theory of creativity (Mednick, 1962), which assumes that creative search is facilitated by semantic memory structure, and with experimental studies linking creativity and semantic network structure (Benedek et al., 2020; Ovando-Tellez et al., 2022) or word associations (Marron et al., 2018). Indeed, the random walks that we compared could be combined into three groups: purely random, structure-driven (frequency-biased), and goal-directed (cue-related adequacy, originality, and likeability biased). We found that the structure-driven random walk outperformed the random and goal-directed random walks, providing further evidence that semantic search has a spontaneous, bottom-up component. Overall, our model is thus compatible with several theoretical accounts of creativity and extends them for instance in terms of phases (generation/evaluation decomposed into exploration, valuation and selection), or in terms of associative theory (showing how spontaneous associations occur during the exploration phase).

### Perspectives

Similar to a previous neuro-computational model of creative processes (Khalil & Moustafa, 2022), our computational model presents the advantage of mathematically formalizing what could be the cognitive operations implemented by the brain during a creative search. This is of importance, as it provides actual variables (such as the likeability of ideas at each trial) that can be related to neural activity, and thus provide insight into the role of each brain region or network involved in the creative process. For instance, the DMN has been identified as a key network for creativity (Beaty et al., 2014), yet it is unclear which computations the different brain regions of this network implement. Our framework, which includes valuation processes, implies that the BVS represents the subjective value of ideas when searching for a creative idea, as this brain network has been found to automatically encode subjective values of any kind of items (Lopez-Persem et al., 2020). Although the BVS has not been frequently reported in previous studies, there is a substantial overlap between the DMN and the BVS, notably in the ventromedial prefrontal cortex and in the posterior cingulate cortex. It is possible that regions considered as belonging to the DMN in previous studies of creativity in fact pertain to the BVS (which deals with idea valuation), while the DMN regions are involved in idea exploration. This hypothetical dissociation has to be directly tested in subsequent studies.

The BVS is also in a good position to interact with the other networks involved in creativity. When making a value-based choice, the BVS interacts with the executive and salience networks in different ways. First, the ECN, including the dorsolateral prefrontal cortex, is thought to regulate – through cognitive control-choices according to the context and goal of the agent (Domenech et al., 2018; Gläscher et al., 2012). For instance, when faced with a food choice between healthy and unhealthy items, the dorsolateral prefrontal cortex (dlPFC, hub of the ECN) has been found to upregulate the weight of the healthy item in the decision (Hare et al., 2009). In our framework, we could speculate that one function of the ECN could be to upregulate the weight of originality in the computation of likeability, to favor more creative outputs and avoid obvious ideas. Second, the salience network, that includes the insula and dorsal anterior cingulate cortex (dACC), is known in neuroscience of decision-making to integrate the decision-value over time to trigger an action selection (Hunt et al., 2014). If the decision-value is close to zero (difficult choice because the two options have close values), the dACC may recruit the dlPFC, to exert some form of control over the choice (better estimating the value of items at stake, for instance) (Shenhav et al., 2013).

Interestingly, the salience network has been proposed to balance the relative involvement of the DMN and ECN in the generation and evaluation processes of creative thinking (Beaty et al., 2016). Some authors have also linked the salience network to a trade-off between exploration and exploitation strategies (Lin & Vartanian, 2018). Thus, the salience network could either play a role in the recruitment of the ECN to exert some control, or to balance the need for exploration (knowledge exploration) and exploitation (maintaining the ongoing idea or strategy) (Kolling et al., 2016). In any case, the value of ideas could be the key missing element in the current framework of creativity. If the value of the current idea is not high enough (low saliency), exploration should be pursued, or re-estimation of value can be performed. Otherwise, the current idea can be further exploited. Future studies will help to specify the role of the dACC and of the salience network in creativity.

In future studies, we will assess the neural bases related to the tasks presented in the current study, and we will focus on the involvement of the BVS and salience network. Additionally, we will assess the generalization of the model with drawings (Barbot, 2018), and Alternative Uses Task (Guilford, 1967). Building networks that could be explored by random walks for those modalities will be challenging, but thanks to the development of various artificial neural networks, similarity matrices (and thus networks) of words (Word2Vec)(Mikolov et al., 2013), concepts (BERT)(Devlin et al., 2019) or drawings (Siamese networks)(Chicco, 2021) can be built. Then, our model will require two inputs: the condition (*First* or *Distant*), mimicking the goal of the participant, stated in the instructions, and the domain (semantic, drawing, or object use). Our framework predicts that only the structure of networks modeling knowledge should differ between modalities, and that valuation and selection functions should be stable across domains.

### Limitations

Some limitations of this study need to be acknowledged. First, the present study assesses creative cognition in the semantic domain. To fully validate our computational model and the core role of preference-based idea selection, it is necessary to apply similar analyses on other domains such as drawings or music. Second, to build our model, we made many assumptions, such as the structure of semantic networks, and each of them should be tested explicitly in future studies. Third, our main result concludes on the role of motivation and preferences in idea selection, but their role in the exploration process per se remains to be further understood.

### Conclusion

The present study reveals the role of individual preferences and decision making in creativity, by decomposing and characterizing the exploration and evaluation/selection processes of idea generation. Our findings demonstrate that the exploration process relied on associative thinking while the selection process depended on the valuation of ideas. We also show how preferences are formed by weighing the adequacy and originality of ideas. By assessing creativity at the group level, beyond the classical interindividual assessment of creative abilities, the current study paves the way to a new framework for creativity research and places creativity as a complex goal-directed behavior driven by reward signals. Future neuroimaging studies will examine the neural validity of our model.

### Data and Code Availability

Data and code will be made available upon publication.

## Acknowledgements

EV is funded by the ‘Agence Nationale de la Recherche’ [grant numbers ANR-19-CE37-001-01]. The research also received funding from the program ‘Investissements d’avenir’ ANR-10-IAIHU-06. MOT is funded by Becas-Chile of ANID. ALP was supported by the «Fondation des Treilles». The study was funded by the Paris Region Fellowship Program, “Horizon 2020 Marie Skłodowska-Curie n° 945298. The overall project to which this study belongs has received funding from the European Union’s Horizon 2020 research and innovation program under the Marie Sklodowska-Curie grant agreement No 101026191. We thank all the participants to the study, Tess Brogard who helped collecting the data, Mehdi Khamassi who provided advices on the model structure, and Bastien Blain for advices on typing speed measures.

