## Supplementary Information for "How Subjective Idea Valuation Energizes and Guides Creative Idea Generation"

This PDF file includes:

Supplementary text

Figures S1 to S9

Tables S1 to S3

SI References

### Supplementary text

#### Results

##### *Choices are predicted by likeability judgements*

We explored whether individuals selected their response in the choice task according to their preferences assessed during the rating tasks. In the choice task, participants had to choose their preferred cue-response association (in the context of the FGAT *Distant* condition) among two options (see Figure 2 and SI Methods *Choice task details*). We examined whether choices were driven by the difference in likeability ratings between the two options (the decision value) (Figure S2). Individual logistic regression of choice rate (in a left-versus-right frame) against decision value (left-minus-right item likeability rating) showed a significant effect at the group level ($\beta_{choice(L)}$=0.04 ± 0.002, t(68)=20.14, p=2.10$^{-30}$). Note that adequacy and originality ratings were also significantly predicting choices ($\beta_{choice(A)}$=0.03 ± 0.002, t(68)=12.77, p=1.10$^{-19}$; $\beta_{choice(O)}$=0.01 ± 0.002, t(68)=3.95, p=2.10$^{-4}$) but the model comparison opposing decision values based on likeability, adequacy and originality ratings revealed that decision value based on likeability better explained choices (Ef=0.74, Xp=1, see SI Methods *Relationship Between Choices and Ratings*).

These results show that when participants are required to make explicit choices for a potential response in the FGAT *Distant* condition, they rely on their subjective values (or likeability).

***Adequacy and Originality ratings can be estimated from associative frequency.***

As likeability ratings are built from adequacy and originality ratings, we checked whether we could predict judgments of adequacy and originality of cue-response associations from their associative frequency (based on *Dictaverf*). We compared several relationships (linear link, quadratic link, both links) between cue-response associative frequency and adequacy and originality, and found that adequacy ratings could be well fitted through a linear relation with frequency ($Ef_{lin}$=0.86, $Xp_{lin}$=1), and that originality could be estimated through a mixture of linear and quadratic link with frequency ($Ef_{lin+quadr}$=0.99, $Xp_{lin+quadr}$=1, Figure S3). Then, for each individual, we had a set of parameters linking adequacy and originality to frequency of a given cue-response association ($\mu_A^l, \mu_O^l\ and\ \mu_O^q$, see Methods in main text: *Valuator Module: Combining Likeability, Originality, and Adequacy of the Rating Tasks with Responses Associative Frequency*). We checked that we could correctly estimate individual ratings of cue-responses adequacy and originality based on their frequency using a leave-one-out procedure on each trial of the rating tasks and for each subject. We found a significant correlation between individual adequacy ratings and estimated adequacy (mean r=0.35±0.01, t(68)=24,48, p=$2.10^{-35}$) and between individual originality ratings and estimated originality (mean r=0.46±0.02, t(68)=20.92, p=$2.10^{-21}$) of cue-responses associations.

### Discussion

#### *Adequacy as the best selection criterion for First Responses*

Our result suggests that adequacy is the best criterion to rely on instead of the first node visited by random walks when selecting the *First* response. While this result might seem counterintuitive at first, it can be easily grasped by considering two non-exclusive aspects. First, semantic networks used in the present study are not individual-based; thus, the first visited nodes might not correspond to the actual individual first associates, and adequacy selection compensates for this approximation. However, adequacy-biased random walks performed worse than frequency-biased random walks; thus, it is also possible that several first associates are triggered by a cue, especially for flat cues, and that a quick adequacy-based decision is made to provide the *First* response. This second interpretation is consistent with the average rank of *First* responses in the fluency task, which was larger than 1 (i.e., 3.24 +/- 0.37), and that suggests that the first associates provided in the FGAT task might belong to a group of first associates. This interpretation is also consistent with the spreading activation model, which assumes that semantic activation spreads from one node (or concept unit) to its neighbouring nodes (Anderson, 1983) possibly in parallel.

### Methods

#### *Experimental Design*

##### FGAT cue words selection and response words

Both conditions used the same list of 62 cue words, presented randomly across each condition. Cue words were nouns, with a lexical frequency (http://www.lexique.org/) above 15 occurrences per million and a number of syllables lower than four. Half of the cues were considered as steep (with a strongly dominant associate) and the other half flat (more balanced strength with its associates). Cues varied in terms of steepness from 1.03 to 24.07. Responses were allowed to be single words (nouns, adjectives or verbs, non-conjugated) avoiding proper nouns and compound words. In each trial, the word given by the participant was automatically searched in a dictionary (http://www.pallier.org/extra/liste.de.mots.francais.frgut.txt) to check for misspelling. If the word was not found in the dictionary at the end of each trial, a window appeared on the screen offering the opportunity to retype the word, and the participant could type in the corrected word or choose to keep their first answer (with no time limit). A secondary phase of manual cleaning cleared the answers from plurals, conjugated verbs, non-words, and typos.

##### Choice task details

Between the likeability rating task and the adequacy-originality rating task, participants performed a binary choice task. They had to choose between two words the one they preferred to be associated with a cue in a creative context, i.e., in the FGAT *Distant* context. Instructions were as follows: ‘For example, would you have preferred to answer “silver” or “jewellery” to “necklace” when generating original associations during the previous task?’ (There was additionally a reminder of the FGAT *Distant* condition, in the instructions).

In each trial, two cue-response associations were displayed on the screen, with the cue word presented on the top of the screen and the two optional responses (the two options of associations) displayed below it on the same horizontal line. Participants had no time limit to make their choice by pressing the left or right arrow on their keyboard. After a 5-

trial training session, participants performed a theoretical maximum number of 519 trials that include the 35 FGAT cues previously selected (or fewer for four participants). Following the cue-word associations built from the rating task (see methods) there was only a maximum of 23 cues with all possible combinations (21*23=483) and a minimum of 12 cues without associations 3,4,5 and 6, thus with 3 possible combinations (3*12)=36, yielding to a total of 483+36=519 theoretical maximal number of combinations. In practice, as cues were presented in the choice task only if they had at least five possible associates, the mean number of choices was 428.6 (median 440) and ranged between 345 and 459, except for three participants who performed 61, 269 and 278 choices.

#### Cue-word associations presented in the rating and choice tasks

The 197 cue-response associations presented in the rating tasks and choice task were built with 35 FGAT cue words randomly selected for each participant, at the end of FGAT with a MatLab script that implemented an adaptive design with the following rules. Each cue word was associated with seven words, amounting to 245 possible associations in total. The seven associated words for each cue word were selected from the participant's answers and from another dataset collected previously in the lab that gathers the responses of 96 independent and healthy participants on a similar FGAT task. They were selected pseudo-randomly with the following constraints that the seven associated words for each given FGAT cue include:

1. A FGAT *First* response given by the participant.
2. A FGAT *Distant* response given by the participant.
3. A frequent FGAT-first response randomly selected among the frequent responses of our independent dataset.

4. An infrequent FGAT *First* response randomly selected among the rare responses of our independent dataset.
5. A frequent FGAT *Distant* response randomly selected among the frequent responses of our independent dataset.
6. An infrequent FGAT *Distant* response randomly selected among the rare responses of our independent dataset.
7. An unrelated word that had no evident semantic link to any of the cue words. We created 35 cue-unrelated word associations that were the same for all subjects.

Note that that association types 3, 4, 5, and 6 were built for only 23 of the 35 FGAT cue words, which were randomly selected for each type, i.e., all associations were created and then associations of type 3 were deleted for twelve random cues, and so on for types 4, 5 and 6 (each time with twelve cues picked randomly among the 35 available cues). This procedure resulted in each cue being associated to three to seven other words. This was done to reduce the number of trials in the rating tasks which was in total 197, except for four participants, in which fewer number of trials were possible (64,141,148 and 169) because of too many missing or misspelled responses in the FGAT.

Frequency of FGAT responses mentioned in associations 3 to 6 corresponds to the number of times an answer was given in the database divided by the total number of answers given for the cue word in our independent dataset. Low and high-frequency responses to a cue are respectively frequencies greater and smaller than the median frequency of responses to a cue.

The associations were in fact pseudo-randomly selected, because we imposed the following constraints to the randomization: for each cue word (i) all seven words had to be different one from another, (ii) composed of at most nine letters, (iii) associated words to a given cue had to be similar in length (with a length difference smaller than five letters), (iv) the three first letters of the seven associates of a cue had to be different, (v) associated words to a given cue and the cue could not "include" each other (e.g. "test" and "retest").

#### Details of the battery of creativity tests

A battery of creativity tests run on Qualtrics followed the previous tasks, in order to assess creative abilities and behavior of the participants. There are described in the order in which they were administered.

##### *Alternative Uses Task (AUT)*

In the AUT task, participants were asked to generate original uses for a common object in three minutes. This procedure was repeated for three objects: tire, bottle and knife. The corresponding nouns naming the objects were presented on the screen during the 3 minutes. Scores for fluency were assessed for each object. Fluency AUT refers to the mean total number of ideas generated by the participant for the three objects.

##### *Inventory of Creative Activities and Achievements (ICAA)*

We used the Inventory of Creative Activities and Achievements (ICAA) questionnaire (64) to assess the real-life creative activities and achievements across eight different creative domains (e.g., literature, music, art and crafts, cooking, sport, visual arts, performing arts, science, and engineering). The creative activities (C-Act) score reflects the frequency in which participants engaged in various creative activities. Six different questions were asked for each domain, and participants reported the frequency with which they engaged in each activity during the last ten years, using an ordinal scale ranging from 0 (never) to 4 (more than ten times). For each participant, the final domain-general score of C-Act was the sum of the creative activities across all activities of the eight different domains. The creative achievements (C-Ach) score estimated the level of achievement acquired in a creative domain. Ten different levels of achievement were included for each domain going from 0 (“never engaged in this domain”) to 10 (“I have already sold some of my work in this domain”). For each participant, the final domain-general score of C-Ach was the sum of the scores across the eight different domains.

##### *Self-report*

Participants were asked to answer the question “How creative would you say you are?” by moving a slider on a continuous scale of 1 to 100, 1 being labeled as “not creative at all” and 100 as “extremely creative”.

##### *Scale of Preferences in Creativity (SPC)*

In the SPC, that we designed in the lab, participants were presented with nine successive questions starting with ‘Would you prefer an idea to be X or Y?’. X and Y were a pair of adjectives with X belonging to semantic field of the word “adequate” and Y belonging to semantic field of the word “original”. The nine pairs were ‘functional’ versus ‘imaginative’; ‘practical’ versus ‘creative’; ‘doable’ versus ‘unique’; ‘efficient’ versus ‘original’; ‘adequate’ versus ‘surprising’; ‘usual’ versus ‘rare’; ‘secure’ versus ‘risky’; ‘certain’ versus ‘ambiguous. Participants had to select a trade-off value on a continuous scale between the two options by moving a slider on a visual scale of -100 to 100, those extremities being labeled as the two written options. The closer the slider was to -100 or 100, the more extremely the participants leaned towards one of the options to the detriment of the other; the closer it was to 0, the more they were ambivalent or showed no preference to either option. The sides of the adequate – original options were counterbalanced. We computed the PrefScore as the mean score across the nine items with more positive scores always indicating preference for originality.

##### *Fluency task*

Participants were presented with a cue word and given 2 minutes to type as many words related to the cue word as possible. The six cue words (‘garden’, ‘wine’, ‘rock’, ‘opinion’, ‘call’ and ‘finger’ - again, note that the cues were written in French for the participants, and the version given here is the closest translation we could find) were selected from the FGAT cues, considering a diversity in steepness.

#### *Statistical Analyses and Computational Modeling*

##### Relationship Between Choices and Ratings

We investigated which rating (Likeability, Adequacy, or Originality) was better explaining choices. Logistic regression was applied to choices as a dependent variable, with likeability (L), originality (O), or adequacy (A) ratings as regressors. Choices were analyzed at the subject level and tested for significance at the group level (random-effect analysis) using two-tailed, paired Student’s t-tests. The softmax function used to determine which variable (V) among likeability, adequacy, or originality ratings was better explaining the proportion choices for left options (P(Left)) against right options is the following:

$$P(Left) = \frac{1}{1 + e^{-\frac{V_{Left} - V_{Right} - d}{\beta_{choice}}}}$$

With *d* being a constant term aiming at capturing any bias towards one side and $\beta_{choice}$ the temperature (choice stochasticity).

#### Construction of Semantic Networks

The *Dictaverf* database consists of 1081 cue words associated with 23340 other words and is organized as a matrix M of i rows and j columns, with associative frequencies directed from the cue-words i to the response-words in j. We used this database combined with FGAT responses to build a symmetric adjacency matrix C of word associations for each cue word, applying the following procedure.

1) A list of all FGAT responses (*First* and *Distant*) from the lab's current and former datasets was created for each cue.
2) Then, for each response in the list:
    a. If it was already associated with the cue in *Dictaverf*, for example, “learning” in response to “school”, then: C(school,learning) = C(learning,school) = M(school,learning).
    b. If it was not associated with the cue in *Dictaverf*: we looked for it in the whole database and identified all potential intermediate nodes between the cue and the word (any other words associated with both the cue and the response). For instance, one subject responded “anxiety” to “school”. “Anxiety” was not directly linked to “school” in M, but it was connected to “studies”, which was connected to “school”. Then, “anxiety” was added as a row (and column for symmetry) in the matrix C, and frequency between “Anxiety” and “studies” was defined as the frequency between “studies” and “anxiety” from M. “Studies” was also added as a row (and column) in C and the frequency between “studies” and “school” was set as the frequency between “school” and “studies”:
    C(anxiety, studies)=C(studies, anxiety)=M(studies, anxiety)

and

C(studies, school)=C(school, studies)=M(studies, school).

In this example, when then building a network based on C (see below), "anxiety" is thus connected to "school" via the node "studies".

This procedure was applied to all potential intermediate nodes, independently of the number of intermediates.

This procedure yielded 62 symmetric C matrices (one per cue) with a size of around 1022 by 1022 words (ranging between 689 and 1186). Each matrix's first row and column correspond to the cue on which the matrix has been built (SI Figure S7 for visual explanation).

Sixty-two networks N were built based on those C matrices as unweighted and undirected graphs (an edge linked two nodes if the frequency of association between them was higher than 0).

##### Random Walks Variants and Implementation

Note that if a given node was not directly linked to the cue, we computed $F_{ci}$ as the cumulative product of the frequency of association of the nodes belonging to the shortest path between the cue and the node. For example: $F_{school\text{-}anxiety} = F_{school\text{-}studies} \times F_{studies\text{-}anxiety}$.

Also, if one node (cue-word association) has actually been rated by the participant, $\mu_A, \mu_O^l, \mu_O^q$, $\alpha$, and $\delta$ were estimated without that particular cue-word association (leave-one-out procedure) to avoid double-dipping. For example, if the cue-response "School-Anxiety" was rated at trial t by a participant, the predicted adequacy, originality, and likeability of trial t for that participant were computed with parameters estimated with all trials except trial t. This procedure lengthens the processing time of the random walks RWA, RWO and RWL but has the advantage of avoiding double-dipping.

#### Decision Functions as the Selector Module

Next, we intended to decipher the criteria determining the selection of a given response. For each subject and cue, we simulated RWF as described above and retained the paths that contained both the *First* and *Distant* response of the subject for further analyses (the number of excluded cues ranged between 0 and 31 trials over 62, M=9.04 trials, exclusion mainly due to missing responses from participants either in the FGAT *First* or *Distant* condition).

For each subject, we built two response matrices $R^F$ and $R^D$ of the same size *n* by *t, t* being the number of cues (equivalent of trials within one FGAT condition) retained (53 cues per subject on average) and *n* the number of nodes visited by the RWF (fixed at 18) (See SI Figure S8 for visual explanation). Those matrices were filled with zeros, except for nodes and trials corresponding to the actual participant response. $R^F$ contains ones for cells actually corresponding to the subject's *First* response (one 1 per column), and $R^D$ contains ones for cells corresponding to the subject's *Distant* response. In order to determine the variable on which the selector module was likely to rely on, we built and compared seven matrices of values $M_x$ of size *n* by *t*:

- $M_r$: random values in the matrix
- $M_P$: matrix with value decreasing with the order in the path
- $M_A$: matrix with estimated adequacy of each visited node in the path
- $M_O$: matrix with estimated originality of each visited node in the path
- $M_L$: matrix with estimated likeability of each visited node in the path

- $M_{A+O:}$ Sum of A and O
- $M_{A*O:}$ Product of A and O

$M_{A+O}$ and $M_{A*O}$ were added as controls for likeability, which relies on a non-linear weighted sum of adequacy and originality (CES).

Using the VBA toolbox, we fitted the following multivariate *softmax* functions to $R^F$ and $R^D$ separately for the seven different matrices:

$$P\left(R_{i,t}^{F}\right) = \frac{e^{-X_{i,t}/\beta^F}}{\sum_{k=1}^{n} e^{(-X_{k,t}/\beta^F)}} \qquad P\left(R_{i,t}^{D}\right) = \frac{e^{-\,(X_{i,t})/\beta^D}}{\sum_{k=1}^{n} e^{(-(X_{k,t})/\beta^D)}}$$

P is the probability of node *i* being selected as a response (R) *First* (F) or *Distant* (D) among all the possible nodes *k* belonging to the *n* options from the paths at trial t (for a given cue). X corresponds to the values within the seven different input matrices. $\beta^F$ and $\beta^D$ are free parameters estimated per subject, corresponding to the temperature (choice stochasticity).

We then compared the seven models for the *First* and *Distant* response separately and reported the results of the model comparison in the results.

### Supplementary figures

**Fig. S1. FGAT Behavior**

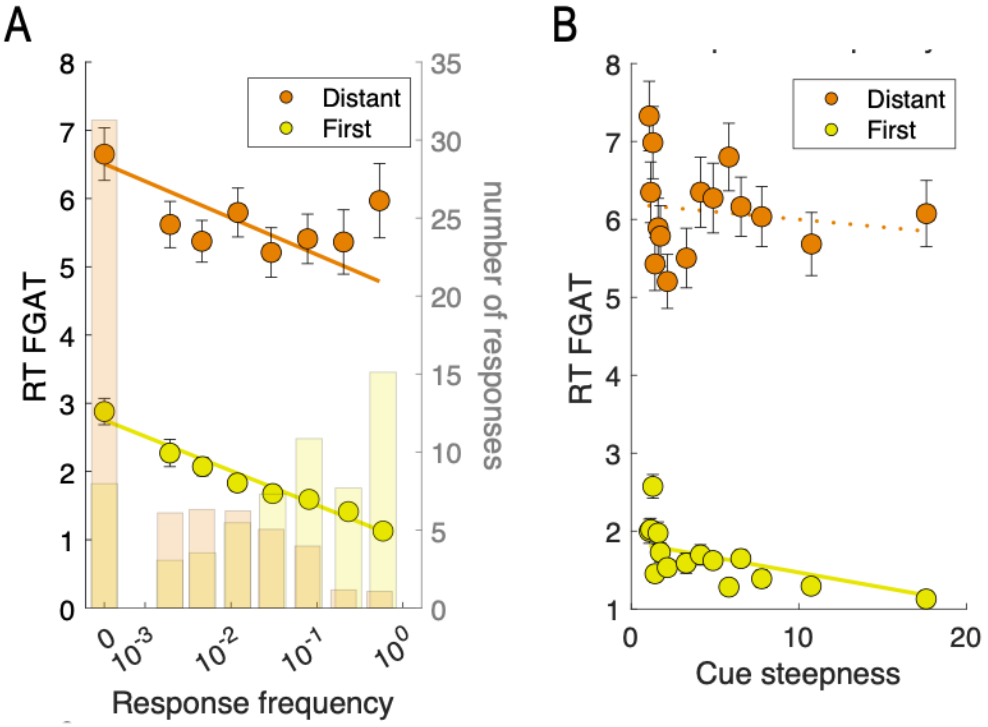


**A.** Correlation between response time (RT) in the FGAT task and the response frequency for the *First* (yellow) and *Distant* (orange) conditions. Transparent bars correspond to the average number of responses per bin of frequency. **B.** Correlation between response time (RT) in the FGAT task and the cue steepness for the *First* (yellow) and *Distant* (orange) conditions. circles indicate binned data averaged across participants. Error bars are intersubject s.e.m. Solid lines correspond to the averaged linear regression fit across participants, significant at the group level ($p<0.05$). Dotted lines indicate that the regression fit is non-significant at the group level ($p>0.05$).

**Fig. S2. Choice task behavior and model fit**

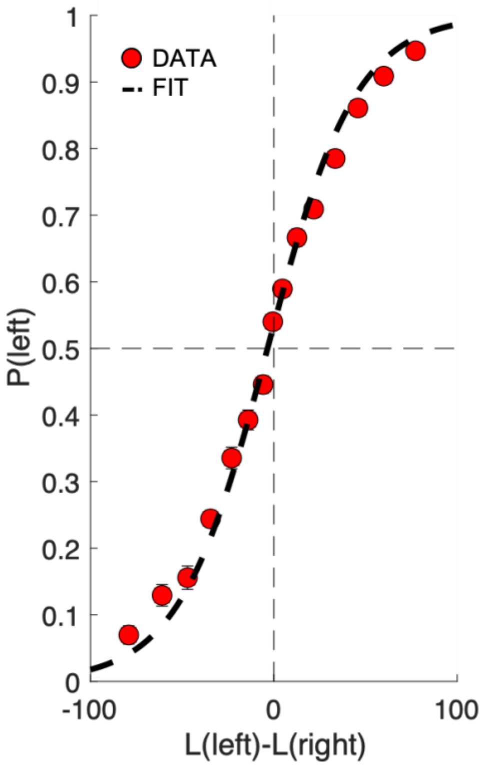


In the choice tasks, the difference between likeability ratings (option values L) predicted choice rate P. Circles indicate binned data averaged across participants. Error bars are intersubject s.e.m. Dashed lines corresponds to the averaged regression fit across participants, significant at the group level ($p<0.05$).

**Fig. S3. Likeability and Frequency of *Distant* Responses in subgroups**

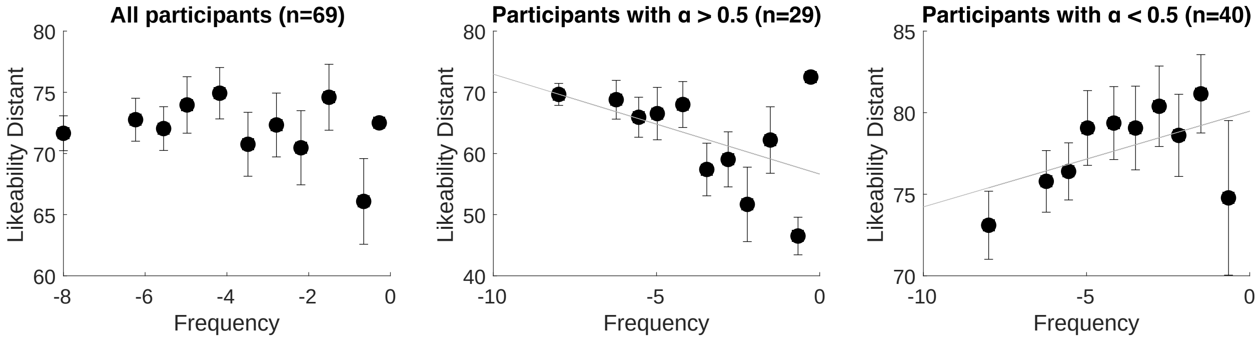


Circles indicate binned data averaged across participants. Error bars are intersubject s.e.m. Solid lines correspond to the averaged linear regression fit across participants, significant at the group level ($p<0.05$).

**Fig. S4. Relationship between association frequency and adequacy/originality ratings**

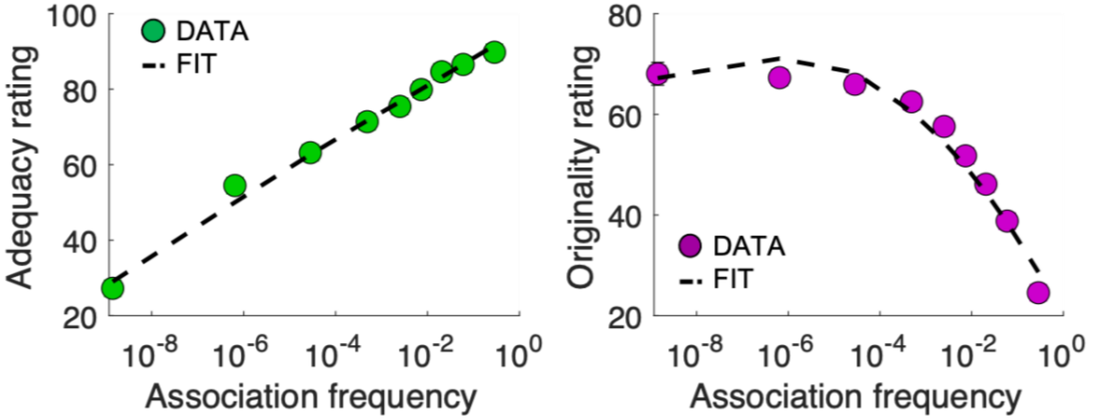


Adequacy (left) and originality (right) ratings in function of frequency of association. Circles indicate binned data averaged across participants. Error bars are intersubject s.e.m. Dashed lines corresponds to the averaged regression fit across participants, significant at the group level ($p<0.05$).

**Fig. S5. Random walks predictions and selection model comparison.**

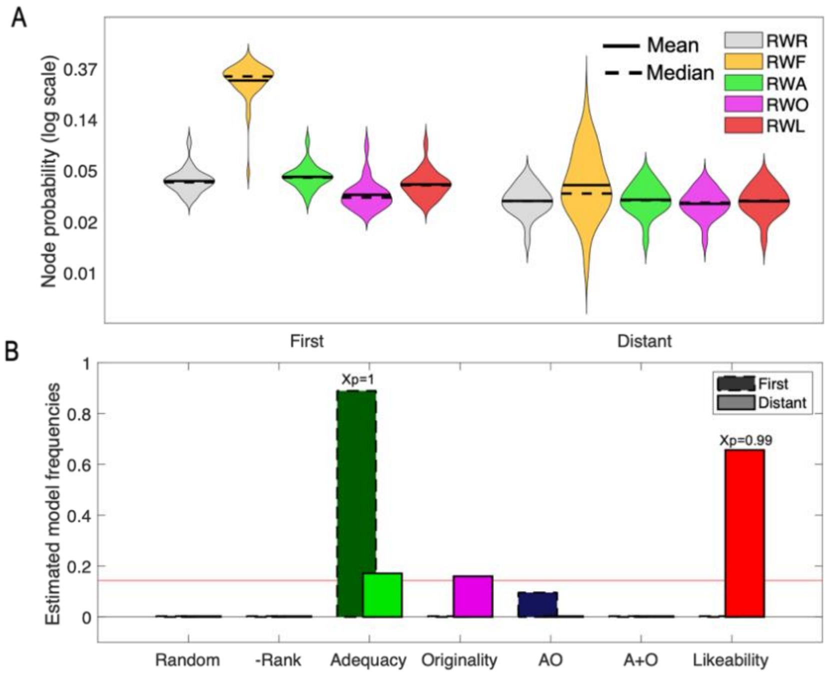


A. Violin plots of the probability of each random walk (RW) to reach the *First* and *Distant* participant responses in the semantic networks. RWR: random, RWF: frequency biased, RWA: adequacy-biased, RWO: originality biased, RWL: likeability biased. Violins represent the distribution of the averaged probabilities across trials for the subgroup of participants used to develop the model (n=46). B. Estimated model frequency of selection models for *First* (dark colors) and *Distant* (lighter colors) responses. Red line indicate chance level.

**Fig. S6. Model and human ranks of responses**

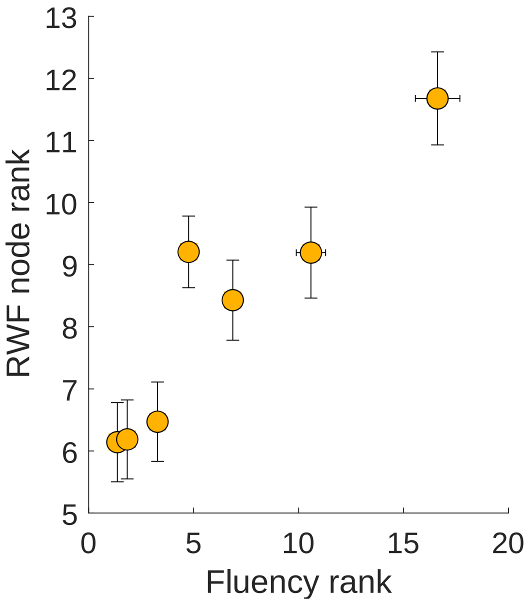


Correlation between the node rank in the RWF model and the fluency rank of words existing both in the fluency responses and model path. Circles indicate binned data averaged across participants. Error bars are intersubject s.e.m.

**Fig. S7. Response quality of the participants and surrogate data of the test group (n=23)**

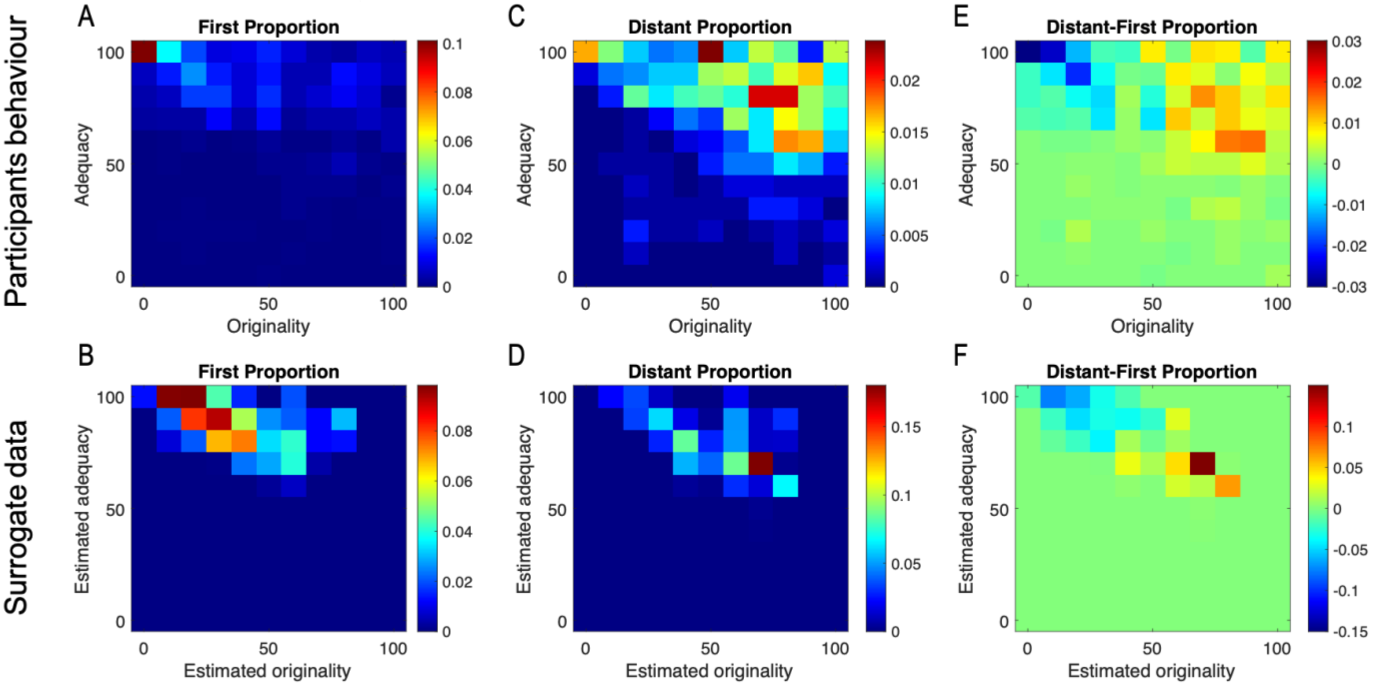


Heatmaps of First (A,B), Distant (C,D) and Distant-First (E,F) proportions of responses per bin of adequacy and originality ratings (A,C,E) or estimated adequacy and originality (B,D,F).

**Fig. S8. Semantic Networks construction**

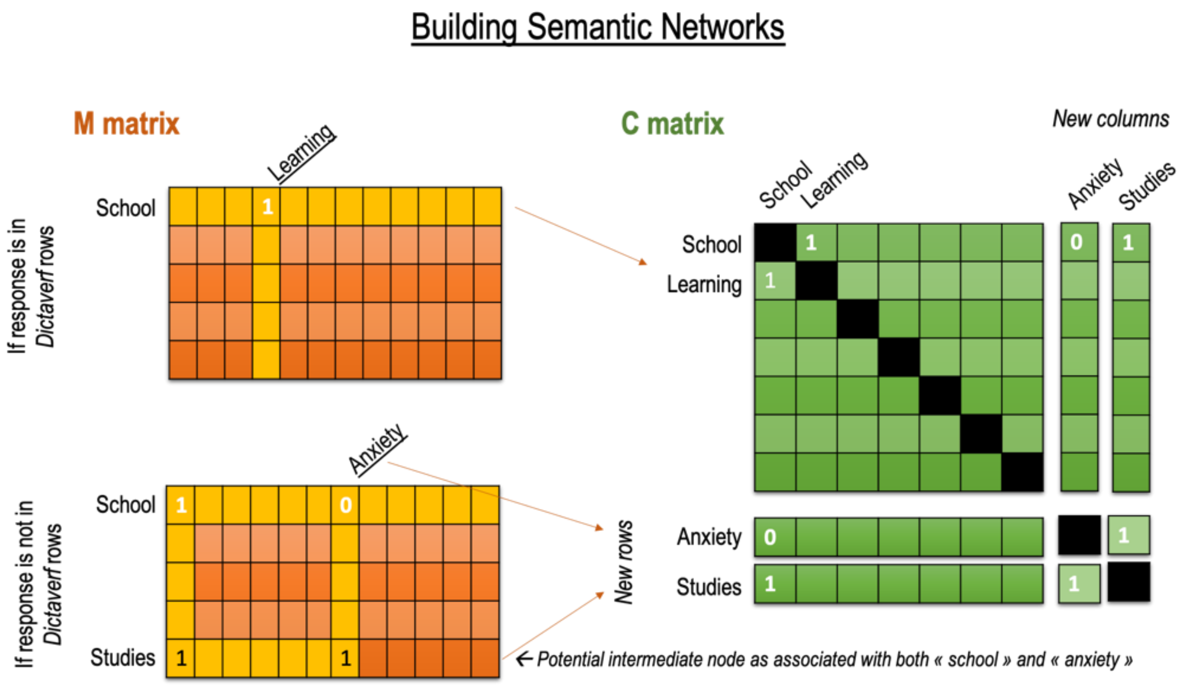


Schematic illustration of the method used to build cue adjacency matrices. See text for details.

**Fig. S9. Illustration of the selector module fitting procedure.**

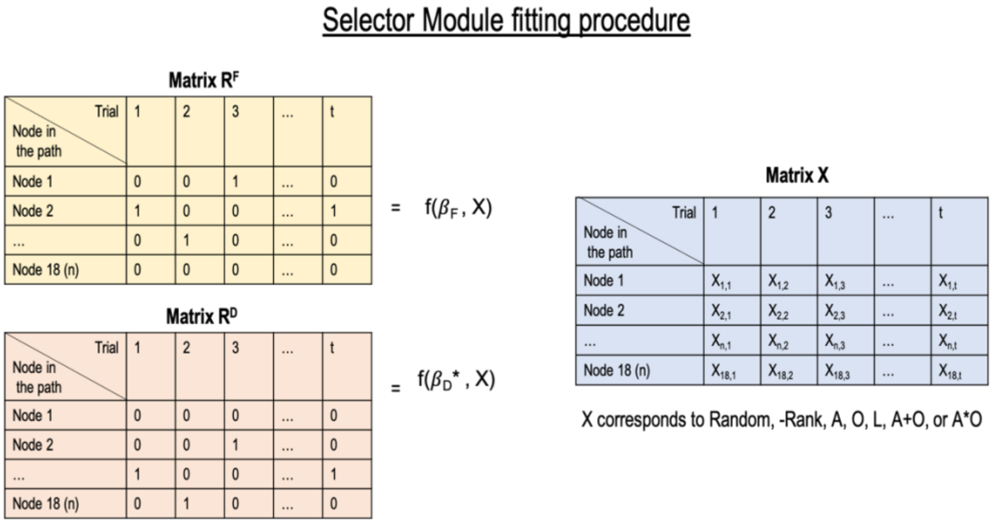


R matrices are the participants responses to be fitted with the selector function f (softmax) against the regressors X that correspond to seven different hypotheses. A: Adequacy, O: originality, L: Likeability. See text for details.

### Supplementary tables

**Table S1. Behavioral results of the FGAT task**

| | | | Mean | SEM | t-value | p-value |
|---|---|---|---|---|---|---|
| Regression of RT against: | Associative frequency | F | -0.34 | 0.02 | -15.92 | $1.10^{-24}$ |
| | | D | -0.10 | 0.02 | -6.27 | $3.10^{-8}$ |
| | | D-F | 0.24 | 0.02 | 10.98 | $1.10^{-16}$ |
| | Steepness | F | -0.13 | 0.02 | -8.50 | $3.10^{-12}$ |
| | | D | -0.02 | 0.01 | -1.16 | 0.25 |
| | | D-F | 0.12 | 0.02 | 5.66 | $3.10^{-7}$ |
| | Likeability | F | 0.08 | 0.02 | 3.78 | $3.10^{-4}$ |
| | | D | -0.15 | 0.02 | -7.25 | $5.10^{-10}$ |
| | | D-F | -0.23 | 0.03 | -7.30 | $4.10^{-10}$ |
| | Likeability (orthogonalized) | F | 0.02 | 0.02 | 1.10 | 0.28 |
| | | D | -0.13 | 0.02 | -6.64 | $6.10^{-9}$ |
| | | D-F | -0.15 | 0.03 | -5.42 | $8.10^{-7}$ |
| Regression of typing speed against: | Likeability | F | 0.01 | 0.02 | 0.36 | 0.72 |
| | | D | 0.08 | 0.02 | 3.88 | $2.10^{-4}$ |
| | | D-F | 0.07 | 0.03 | 2.21 | $3.10^{-2}$ |
| | Likeability (orthogonalized) | F | 0.02 | 0.02 | 0.90 | 0.37 |
| | | D | 0.07 | 0.02 | 3.16 | $2.10^{-3}$ |
| | | D-F | 0.05 | 0.03 | 1.54 | 0.13 |

**Table S2. Statistical results about the human behavior of the test group and the surrogate data.**

| | | | Human behavior | | | | Surrogate data (100 simulations per individual) | | | |
|---|---|---|---|---|---|---|---|---|---|---|
| | | | Mean | SEM | t-value | p-value | Mean | SEM | t-value | p-value |
| Regression of RT/Rank against: | Group frequency | F | -0.27 | 0.04 | -6.98 | $5.10^{-7}$ | -0.14 | 0.00 | -53.75 | $8.10^{-25}$ |
| | | D | -0.11 | 0.03 | -3.70 | $1.10^{-3}$ | -0.05 | 0.01 | -4.35 | $3.10^{-4}$ |
| | | D-F | 0.16 | 0.04 | 4.04 | $5.10^{-4}$ | 0.09 | 0.01 | 6.43 | $2.10^{-6}$ |
| | Steepness | F | -0.11 | 0.03 | -4.09 | $5.10^{-4}$ | -0.09 | 0.00 | -43.27 | $9.10^{-23}$ |
| | | D | -0.02 | 0.02 | -0.89 | 0.381 | -0.03 | 0.01 | -2.42 | 0.024 |
| | | D-F | 0.09 | 0.03 | 2.60 | 0.016 | 0.07 | 0.01 | 6.07 | $4.10^{-6}$ |
| | Likeability | F | 0.04 | 0.03 | 1.37 | 0.185 | 0.05 | 0.04 | 1.45 | 0.162 |
| | | D | -0.17 | 0.05 | -3.73 | $1.10^{-3}$ | -0.05 | 0.02 | -2.20 | 0.039 |
| | | D-F | -0.21 | 0.06 | -3.54 | $2.10^{-3}$ | -0.10 | 0.02 | -4.91 | $6.10^{-5}$ |
| Mean effects | Adequacy | F-D | 9.24 | 0.99 | 9.29 | $1.10^{-13}$ | 10.24 | 2.56 | 4.00 | $6.10^{-4}$ |
| | Originality | D-F | 30.63 | 1.87 | 16.36 | $3.10^{-25}$ | 17.55 | 4.17 | 4.21 | $4.10^{-4}$ |
| | Originality-Adequacy | D-F | 21.40 | 1.54 | 13.87 | $2.10^{-21}$ | 7.31 | 2.08 | 3.51 | $2.10^{-3}$ |

F stands for *First* responses; D stands for *Distant* responses.

**Table S3. Coefficient of correlation between each variable and its corresponding canonical variable**

| Sets | Variables | coefficients | p-values |
|---|---|---|---|
| FGAT individual parameters & scores | α (weight A/O) | 0.38 | 0.001 |
| | δ (convexity A/O) | 0.54 | $2.10^{-6}$ |
| | $\beta_{choice}$ (Choice temperature) | 0.02 | 0.88 |
| | $\beta_{selection}$ (*Distant* temperature in *selector* module) | 0.26 | 0.03 |
| | -Associative Frequency cue-*First* | 0.76 | $2.10^{-14}$ |
| | -Associative Frequency cue-*Distant* | 0.15 | 0.22 |
| Battery individual scores | AUT fluency | 0.37 | 0.002 |
| | Fluency | 0.29 | 0.017 |
| | C-Act | 0.07 | 0.55 |
| | C-Ach | -0.16 | 0.18 |
| | Creativity Self-report | 0.48 | $4.10^{-5}$ |
| | PrefScore | 0.60 | $5.10^{-8}$ |
